# An antibiotic that mediates immune destruction of senescent cancer cells

**DOI:** 10.1101/2024.09.05.611241

**Authors:** Gabriele Casagrande Raffi, Jian Chen, Xuezhao Feng, Zhen Chen, Cor Lieftink, Shuang Deng, Jinzhe Mo, Chuting Zeng, Marit Steur, Jing Wang, Onno B. Bleijerveld, Liesbeth Hoekman, Nicole van der Wel, Feng Wang, Roderick Beijersbergen, Jian Zheng, Rene Bernards, Liqin Wang

## Abstract

Drugs that eliminate senescent cells, senolytics, can be powerful when combined with pro-senescence cancer therapies. Using a CRISPR/Cas9-based genetic screen, we identify here SLC25A23 as a vulnerability of senescent cancer cells. Suppressing SLC25A23 disrupts cellular calcium homeostasis, impairs oxidative phosphorylation and interferes with redox signaling, leading to death of senescent cells. These effects can be replicated by salinomycin, a cation ionophore antibiotic. Salinomycin prompts a PANoptosis-like cell death in senescent cells, including apoptosis and two forms of immunogenic cell death: necroptosis and pyroptosis. Notably, we observed that salinomycin treatment or SLC25A23 suppression elevates reactive oxygen species, upregulating death receptor 5 via JNK pathway activation. We show that a combination of a DR5 agonistic antibody and salinomycin is a robust senolytic cocktail. We provide evidence that this drug combination provokes a potent NK and CD8+ T cell mediated immune destruction of senescent cancer cells, mediated by the pyroptotic cytokine IL18.

**Significance:** The efficacy of multiple cancer drugs is limited by the induction of senescence, which enables cancer cells to evade cell death. We uncover here a new selective vulnerability of senescent cancer cells and we show that this vulnerability can be targeted with a commonly-used antibiotic. We show that a combination of senescence-inducing therapy, combined with this antibiotic causes a highly immunogenic cell death phenotype that further stimulates potent immune destruction of senescent cancer cells.

## Introduction

Senescence is a stable proliferation arrest that can result from various stresses and which serves as a fail-safe mechanism against the development of cancer.^1^ Senescence can also be induced in cancer cells, for instance, as a result of genotoxic stress brought about by chemotherapeutic agents.^2^ Despite the apparent advantage of proliferation arrest in cancer cells, studies have highlighted that the various cytokines produced by senescent cancer cells, collectively referred as the senescence-associated secretory phenotype (SASP), can be a double-edged sword for tumor control.^3^ On the one hand, the SASP suppresses tumor growth by inducing an immunological response. On the other hand, prolonged tumor exposure to SASP can be harmful, causing chronic inflammation, immunosuppression, epithelial-to-mesenchymal transition, stem cell-like state induction, or tumor migration and metastasis promotion.^4^ Therefore, while senescence-induced growth arrest may be a desirable outcome initially, the prolonged presence of senescent cells can have detrimental consequences.

The intricate interplay between senescent cancer cells and the immune microenvironment remains a topic of ongoing investigation, with many aspects yet to be fully elucidated. The release of Damage-Associated Molecular Patterns (DAMPs) upon senescent cell clearance has the potential to activate an immune response, creating a pro-inflammatory environment.^5^ Indeed, evidence has been presented that induction of senescence sensitizes pancreatic cancer cells to anti-PD-1 therapy.^6^ Conversely, the release of SASP factors can also exert an immunosuppressive effect, dampening the immune response.^7,8^ This raises questions regarding the context in which senescent cells can modulate the inflammatory phenotype associated with these cells.^9^

Senolytics, agents that kill senescent cells, have emerged as a promising therapeutic strategy to selectively eliminate senescent cells that accumulate in tissues during aging and in age-related diseases, including cancer.^2,10^ Eliminating these cells is believed to improve tissue function and delay or reverse the aging process.^11,12^. In cancer therapy, senolytics could be combined with pro-senescence therapies to prevent the harmful effects of persistent senescent cells in tumors and during cancer treatment.^13^ Additionally, most of the therapies developed in the clinic mainly target actively proliferating cells, and a state associated to the withdrawal from cell cycle, such as senescence, can render these therapies ineffective. This highlights the clinical need to combine senescence inducing therapies, namely chemotherapy or radiotherapy, with senolytics to effectively eradicate both proliferating and non-proliferating cells. Notwithstanding the potential benefits of this strategy, the development of senolytic drugs has encountered obstacles due to the absence of universal agents that can effectively target senescent cancer cells in a manner that is not dependent on the cellular context while maintaining an acceptable toxicity profile.^14,15^ Identifying more universal vulnerabilities in senescent cancer cells would represent a significant step forward in developing effective senolytic therapies. As more targets are identified and new drugs are developed, senolytic therapy could potentially become an approach to combine with senescence-inducing therapies for treating cancer.^2,10^

Here, we have used unbiased genetic screens to identify targetable vulnerabilities in senescent cancer cells. These screens involve the systematic loss-of-function analysis of genes essential for senescent cells’ survival but not for proliferating cancer cells. We find an unexpected new vulnerability of senescent cancer cells that can be targeted with a commonly used antibiotic drug to kill senescent cancer cells.

## Results

### CRISPR-based senolytic screen identifies SLC25A23 as a senolytic target

To identify genes whose inactivation causes cell death selectively in senescent cancer cells, we re-analyzed a previously performed genome-scale loss of function genetic screen using a doxycycline-inducible CRISPR/Cas9 vector system.^16^ In brief, we used as the screening model A549 KRAS mutant lung cancer cells treated with an aurora kinase A inhibitor alisertib to induce senescence. We first introduced the lentiviral gRNA library in proliferating A549 cells, followed by selection for lentiviral integration. The gRNA-expressing cells were treated with alisertib to induce senescence. After this, CAS9 expression was activated by doxycycline treatment to induce gene knock-outs. The senescent cell populations were cultured for 10 days to allow for the depletion of gRNAs in the senescent cells (Fig. 1a). A control arm of proliferating cells was also included in the screen. We selected sgRNAs preferentially depleted from cells made senescent by alisertib, with little or no effect in the proliferating arms. Using this criterium, we have previously identified cFLIP as a vulnerability of senescent cancer cells.^16^ Further data analysis identified another significantly depleted gene in senescent A549 cells: SLC25A23 (Fig. 1b). SLC25A23 is a mitochondrial inner membrane protein known for regulating mitochondrial calcium homeostasis. Other reports highlight how calcium homeostasis in mitochondria can control key functions like mitochondrial protein translation, respiratory chain functions and maintaining redox balance^17–20^. We validated the hit using two independent gRNAs targeting SLC25A23 in A549 cells harboring the doxycycline-inducible Cas9 (A549-iCas9). Treatment of these cells with doxycycline significantly reduced the SLC25A23 mRNA expression (AI Appendix, Fig. S1a). Colony formation assays confirmed that reducing SLC25A23 expression can preferentially kill the senescent A549 cells (AI Appendix, Fig. S1b-c). We also used three independent shRNAs targeting SLC25A23 in A549 cells to confirm this phenotype further. SLC25A23 suppression was measured through RT-qPCR (Fig. 1c). Loss of SLC25A23 induced cell death not only in alisertib-induced senescent cells but also in cells made senescent by two other agents: the PLK4 inhibitor CFI-400945 and the chemotherapeutic agent etoposide (Fig. 1d). Efficient induction of senescence was verified using staining for the senescence marker senescence-associated β-galactosidase for all the models (Supplementary Fig. S1d). Senolytic activity was also observed upon SLC25A23 suppression in SK-Hep1 liver cancer cells and MDA-MB-231 breast cancer cells (AI Appendix, Fig. S1e-h).

**Figure 1.**
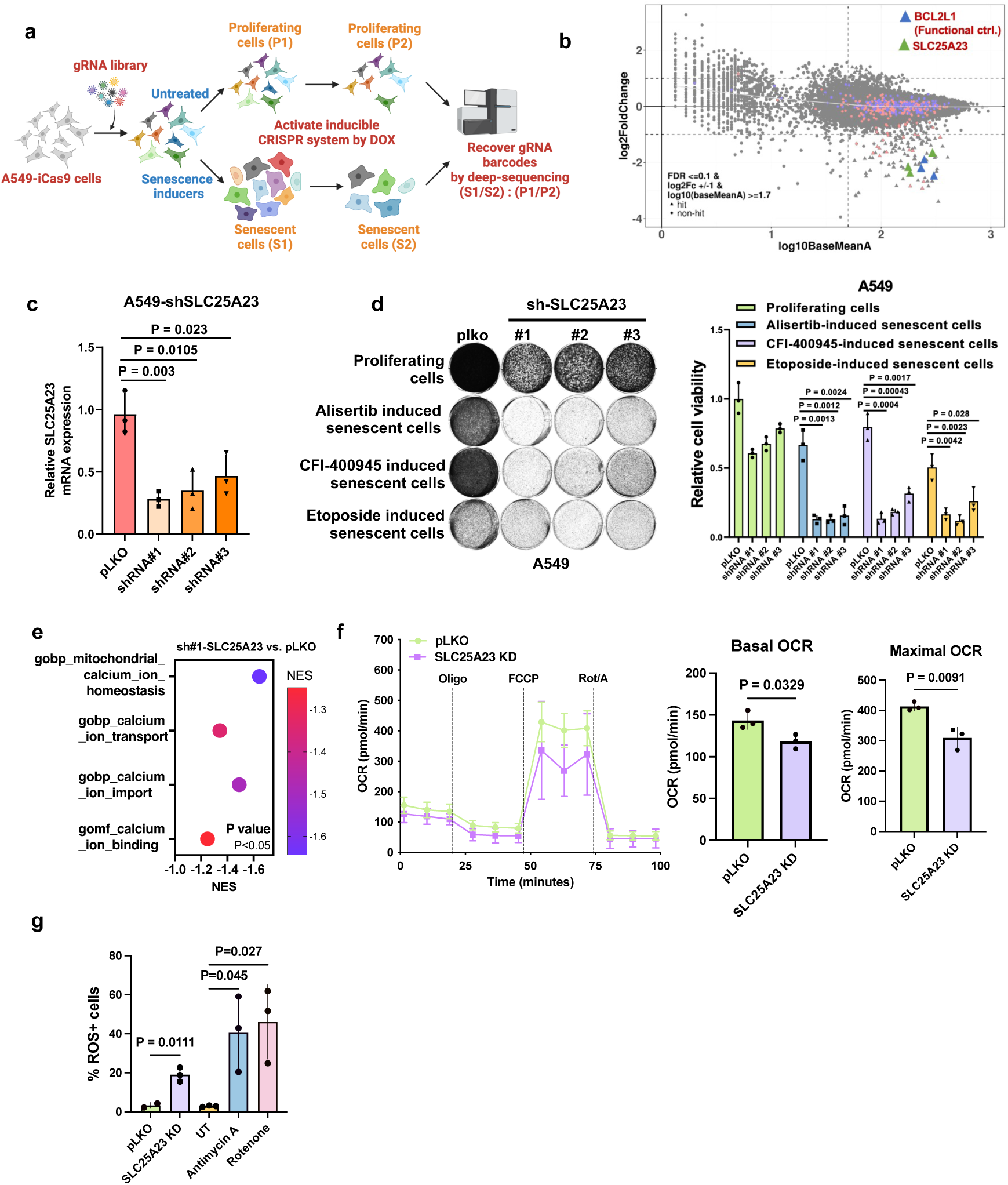
CRISPR-based senolytic screen identifies SLC25A23 as a senolytic target. (a). Schematic of iCas9-associated senolytic CRISPR screen workflow. Brunello lentiviral whole genome-wide gRNA collection virus was introduced to A549 cells infected with a doxycycline-inducible Cas9 (iCas9) lentiviral vector. Alisertib-induced senescent cells and proliferating cells were treated with 1 µg ml^-1^ doxycycline (DOX) for 10 days, and Illumina deep-sequencing was used to determine changes in library representation. (b). gRNAs prioritized for further analysis were selected by the fold depletion of abundance in the S2 and S1 samples compared with that in P2 and P1 samples. SLC25A23 was identified as the top hit. BCL2L1 was considered a positive control. (c). To validate the knockdown efficiency, we performed Real-time PCR analysis of SLC25A23 in A549-iCas9 cells with three independent shRNAs against SLC25A23. GAPDH served as a loading control. n = 3 independent experiments. Error bars represent the mean ± standard deviation. One-Way ANOVA was performed to calculate statistical significance. (d). Cell viability of proliferating and alisertib-, CFI-400945- and etoposide-induced senescent A549 cells with three independent shRNAs against SLC25A23 was assessed by Colony Formation Assay. pLKO empty vectors were taken as a control. Cell viability is quantified. n = 3 independent experiments. Error bars represent the mean ± standard deviation. Two-Way ANOVA was performed to calculate statistical significance. (e). Bobble plot showing pathway enrichment of mitochondrial calcium ion homeostasis, calcium ion transport, calcium ion imports and calcium ion binding in shRNA SLC25A23 compared to pLKO empty vector A549 cells. (f). Oxygen consumption rate (OCR) in A549 cells with shRNA against SLC25A23 was measured by Seahorse assay. pLKO empty vectors were taken as a control. Basal OCR and maximal respiratory capacity were quantified. For both n = 3 time points, and error bars represent mean ± standard deviation. Two-tailed unpaired student’s t-test was performed to calculate statistical significance. (g). Flow cytometry was used to determine the percentage of ROS-positive cells in A549 cells with shRNA against SLC25A23 and in A549 cells treated with 2 µM Antimycin A or 2 µM Rotenone for 24 hours. pLKO empty vector and untreated A549 cells were taken as a control, respectively. n = 3 independent replicates. Error bars represent mean ± standard deviation. Two-Way ANOVA was performed to calculate statistical significance.

SLC25A23 is involved in multiple biological functions, which may be critical for the survival of senescent cells.^21–24^ The suppression of SLC25A23 decreases mitochondrial calcium, leading to cytosolic calcium accumulation and deregulated cellular calcium homeostasis, affecting mitochondrial Electron Transport Chain (ETC) complex assembly, redox homeostasis and inducing Endoplasmatic Reticulum (ER) instability.^17,20,25,26^ We observed that SLC25A23 suppression could induce a decrease in calcium signaling through RNA-seq measurements, with some signatures describing calcium homeostasis and calcium signaling being downregulated (Fig. 1e). We also detected an increase in cytosolic calcium as measured using flow cytometry judged by the intensity of the calcium-specific dye Rhod-2-AM, highlighting the role of SLC25A23 in controlling mitochondrial and cellular calcium balance. We then tested whether this calcium imbalance induced by the downregulation of SLC25A23 in A549 cells could impair mitochondrial activity and OXPHOS efficiency by measuring oxygen consumption rate (OCR) using Seahorse analysis. We observed a significant reduction in the basal OCR and the maximal respiratory capacity in the SLC25A23 knock-down cells (Fig. 1f). Consequently, we observed an increase in cellular Reactive Oxygen Species (ROS) induced by the suppression of SLC25A23, similar to standard OXPHOS inhibitors (Fig. 1g). These results highlight that SLC25A23 is important in controlling the cell’s calcium homeostasis, affecting mitochondrial activity, and potentially inducing ER stress.^26^

### Salinomycin phenocopies SLC25A23 suppression-mediated senolysis

SLC25A23 depletion induces senolysis in multiple tumor models rendered senescent using different senescence inducers. However, to date no known direct inhibitors of the protein have been identified. To find a drug that could phenocopy the biological effects of SLC25A23 suppression, we performed a senolytic drug screen with a mini-library of drugs selected to act on cellular processes regulated by SLC25A23, including calcium homeostasis, mitochondrial functions and OXPHOS, and ER stress (Table S1). We included navitoclax (ABT-263) as positive control for senolysis. The results of this drug screen are represented according to each drug’s senolytic index log_2_ (IC50PAR/IC50SEN). As the top hit of the screen, the cation ionophore antibiotic salinomycin was identified (Fig. 2a). We then validated the senolytic activity of salinomycin in different models. These experiments showed that salinomycin is capable of selectively killing the senescent cells over proliferating cells in multiple cell models, including lung (A549), breast (MDA-MB-231, MCF-7, T47D), liver (SK-Hep1) and melanoma (SK-Mel28), which were made senescent using different senescent inducers (Fig. 2b and AI Appendix Fig. S2a-c). We conducted proteomic and RNA-seq analyses to investigate the impacts of salinomycin on cellular physiology. Analysis of the RNA-seq data revealed a significant reduction in the activity of crucial and non-redundant calcium signaling pathways (Fig. 2c and AI Appendix Fig. S2d), decreased OXPHOS activity (Fig. 2c, d) and increased unfolded protein response (UPR) (Fig. 2c, d). These observations were further confirmed through functional assays. We detected an increase in cytosolic calcium levels through flow cytometry measurement of the calcium-specific dye Rhod-2 AM, indicating deregulation of calcium homeostasis (Fig. 2e). We also noticed a reduction in OXPHOS efficiency of the salinomycin-treated cells, evident in both basal OCR and maximal respiratory capacity (Fig. 2f). Consequently, we observed increased basal ROS levels in the salinomycin-treated cells in both parental and senescent states (Fig. 2g). We treated senescent A549 cells with salinomycin with and without N-acetyl cysteine (NAC), the most widely used ROS scavenger, to see whether ROS play a key role in killing the senescent cells. We observed that NAC could not reduce salinomycin’s killing capacity over a range of concentrations (AI Appendix Fig. S2e). In summary, salinomycin affects calcium homeostasis, inducing ER stress, mitochondrial dysfunction and an increase in cellular ROS levels, similar to what previously observed with SLC25A23 depletion. These data suggest that despite the lack of a direct SLC25A23 inhibitor, salinomycin phenocopies the loss of SLC25A23 and is capable of inducing strong senolytic effect in multiple models.

**Figure 2.**
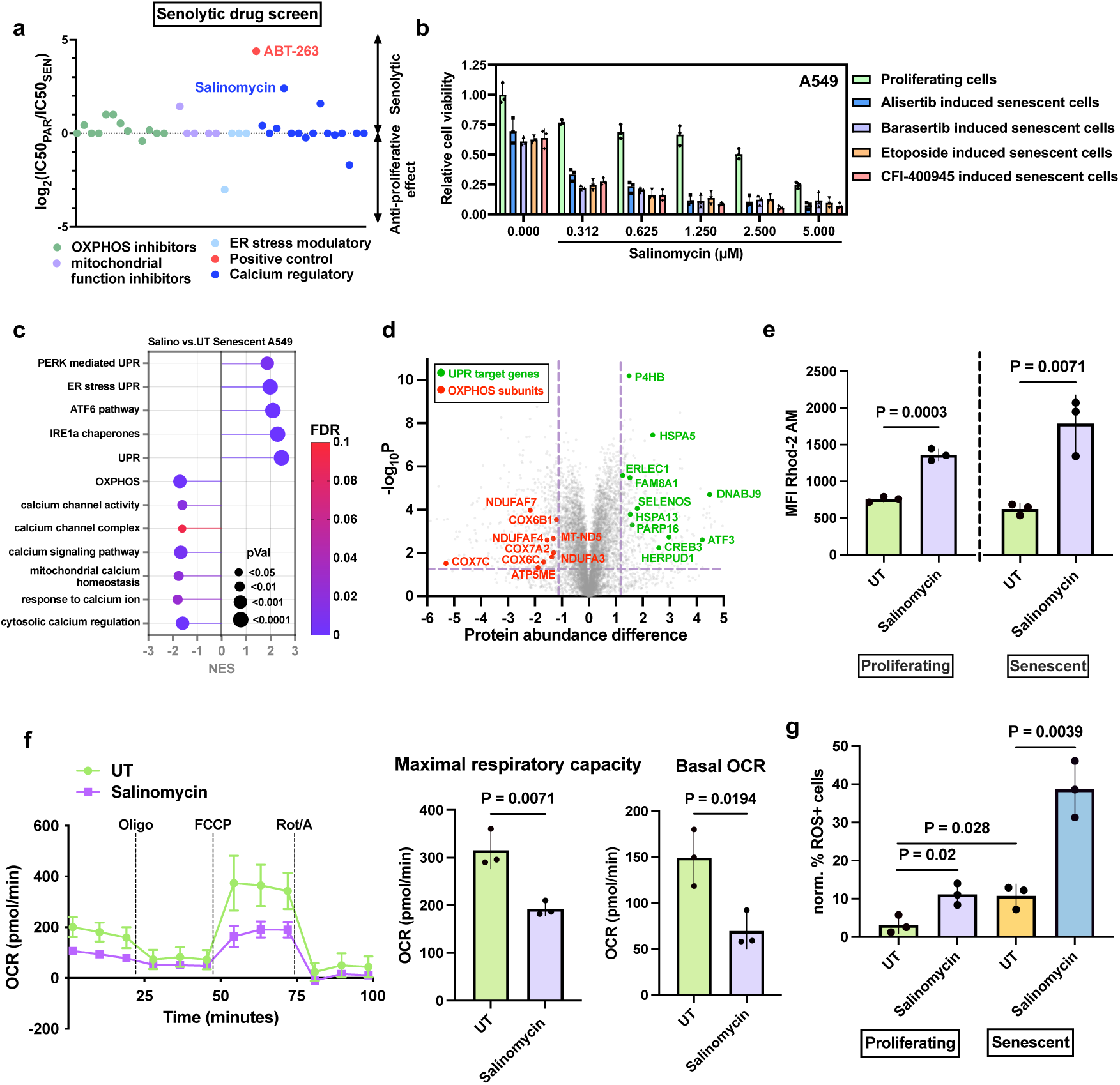
Salinomycin phenocopies SLC25A23 depletion and induces senolysis. (a). Senolytic indexes of OXPHOS and mitochondrial complex inhibitors, mitochondrial function inhibitors, ER stress modulatory drugs and calcium regulatory drugs are determined by dividing the IC50 value of the drugs in parental A549 cells by the IC50 value in alisertib-induced senescent A549 cells. (b). Cell viability of alisertib-, barasertib-, etoposide- and CFI-400945-induced senescent A549 cells treated with salinomycin was assessed by colony formation assay. Proliferating cells were taken as a control. The cell viability was quantified (right panel). n = 3 independent experiments. Error bars represent mean ± standard deviation. Statistical significance was calculated with Two-Way ANOVA but is not shown in the graph. (c). Bobble plot showing pathway enrichment of calcium channel activity, calcium channel complex, calcium signaling pathway, mitochondrial complex homeostasis, OXPHOS, PERK, IRE1a, ATF6, UPR response to calcium ion and cytosolic calcium regulation in alisertib-induced senescent A549 treated with 2.5 µM salinomycin for 48 hours compared to untreated cells. The size of the circle indicates the p-value. (d). Volcano plot showing differentially expressed proteins related to OXPHOS subunits (red) and Unfolded Protein Response and ER stress (green) in alisertib-induced senescent A549 cells treated with 2.5 µM salinomycin for 48 hours compared to untreated cells. Thresholds for significance are -1.2 and 1.2 for abundance difference and 1.3 for -log_10_P. (e). Flow cytometry was used to determine the intensity of Ca^2+^ dye Rhod-2 in parental and alisertib induced senescent A549 cells treated with 2.5 µM salinomycin for 4 hours. Untreated cells were taken as a control. n = 3 independent experiments. Error bars represent the mean ± standard deviation. Two-tailed unpaired student’s t-test was performed to calculate statistical significance. (f). Oxygen consumption rate (OCR) in proliferating A549 treated after 24 hours of 2.5 µM salinomycin treatment was measured using Seahorse assay. Untreated cells were taken as a control. Basal OCR and maximal respiratory capacity were quantified. For both n = 3 independent experiments, error bars represent mean ± standard deviation. Two-tailed unpaired student’s t-test was performed to calculate statistical significance. (g). Flow cytometry determined the percentage of ROS-positive cells in proliferating and alisertib-induced senescent A549 cells after 24 hours of 2.5 µM salinomycin treatment. Untreated proliferating cells were taken as a control. n = 3 independent experiments. Error bars represent the mean ± standard deviation. One-way ANOVA was performed to calculate statistical significance.

### Salinomycin induces PANoptosis-like cell death in senescent cells and upregulates DR5

Next, we studied how salinomycin induces cell death. Under the microscope, we observed that senescent cells treated with salinomycin undergo an unusual cell death morphology (Supplementary Fig. S3a).^27^ This observation was confirmed through Scanning Electron Microscope images (Fig. 3a). We found through western blot analysis that multiple death pathways are active in parallel. Our analyses revealed heightened levels of both total and phosphorylated MLKL, a pivotal mediator in necroptotic demise (Fig. 3b). Furthermore, the observed elevation in cleavage of caspase 1 (cl-CASP1) and cleaved GSDMD (cl-GSDMD) provides evidence for pyroptosis induction (Fig. 3b). Moreover, our data demonstrate salinomycin’s induction of cleaved forms of caspase 8 and caspase 3, alongside a noteworthy upregulation of DR5, collectively implicating activation of the extrinsic apoptotic cascade (Fig. 3b). These observations were validated in the SK-Hep1 cell line (AI appendix Fig. S3b) and the *SLC25A23* knockdown cells treated with alisertib (AI appendix Fig. S3c), further confirming that salinomycin can phenocopy the behavior of *SLC25A23* suppression with respect to the types of death. We also confirmed this phenotype by studying the RNA-seq data. We found that salinomycin modulates the mRNA expression of key factors involved in necroptosis (Fig. 3c). Notably, our data also suggest that senescent cells are primed for pyroptotic death. This is supported by the notion that senescent cells upregulate the basal levels of pyroptosis gene signature (AI appendix Fig. S3d), which is consistent with the observations showing an increase in the basal levels of cleaved gasdermin D (cl-GSDMD), the pore-forming molecule involved in pyroptotic death (Fig. 3b). This is also confirmed by the finding that multiple senescent cell lines are more sensitive to the pyroptosis inducer nigericin than their parental counterpart (AI appendix Fig. S3e). The induction of pyroptosis was further validated by the release in the supernatant of inflammatory molecules like IL-18 and IL1b by the salinomycin-treated senescent cells (Fig. 3d, e, AI appendix Fig. S3f). These results suggested that pyroptosis was primed during senescence induction and further stimulated by salinomycin. Our collective evidence suggests that salinomycin induces not only singular, but concurrent activation of multiple cell death pathways. To verify that multiple cell death mechanisms contribute to the senolytic effect of salinomycin, we used multiple death inhibitors to see which ones could rescue salinomycin-induced death. We used selected single, double and triple combinations of the death inhibitors Q-VD-OPh (apoptosis), Z-VAD-FMK (apoptosis), necrosulfonamide (necroptosis) and MCC-950 (pyroptosis) to observe the contribution of each of these pathways to senescent cells death. We observed that no inhibitor as a single agent could block salinomycin-induced killing; double combinations could partially prevent salinomycin-induced killing, but only with a three-way combination could we achieve full death rescue (Fig. 3f). These results strongly indicate that salinomycin mediates cell death through multiple mechanisms acting in parallel.

**Figure 3.**
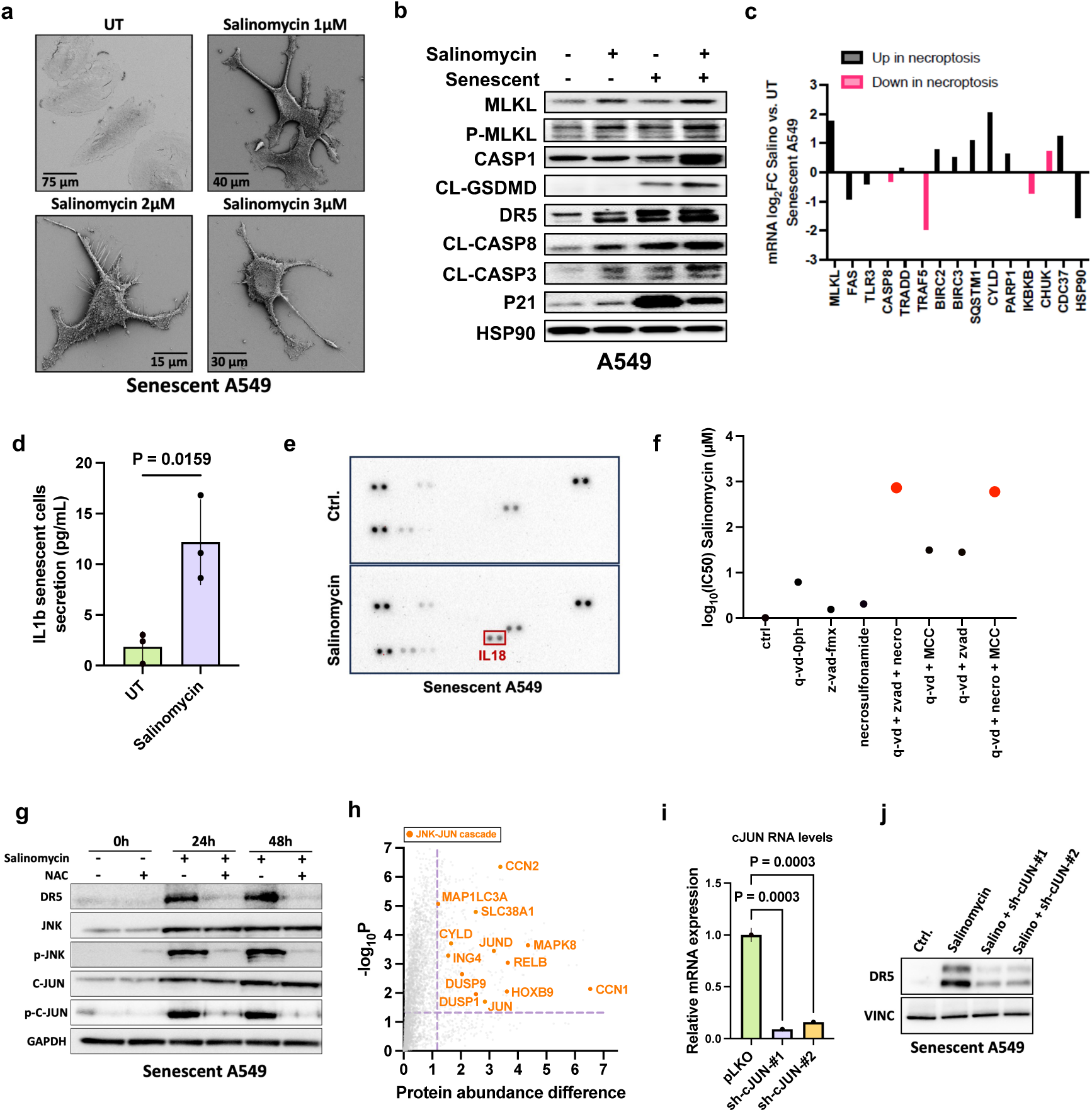
Salinomycin induces cell death through multiple parallel mechanisms. (a). Images taken with scanning electron microscopy show the morphology of alisertib-induced senescent A549 cells after 3 days of 1 µM, 2 µM or 3 µM salinomycin treatment. Untreated alisertib-induced senescent cells were taken as a control. (b). Western blot analysis of apoptosis, pyroptosis, necroptosis death pathway markers in alisertib-induced senescent and proliferating A549 cells after 48 hours of 2 µM salinomycin treatment. HSP90 served as loading control. (c). Bar plot showing differentially expressed necroptosis-related genes in alisertib-induced senescent A549 cells after 48 hours of 2.5 µM salinomycin treatment compared to untreated cells. mRNA expression was measured by bulk RNA-sequencing. The bars represent the average value of 3 independent replicates. The plot shows the genes upregulated (black) and downregulated (red) upon necroptosis induction as a function of log_2_FC of salinomycin-treated cells versus untreated (UT). (d). IL1β release of alisertib-induced senescent A549 cells after 48 hours of 2.5 µM salinomycin treatment was measured using IL1β Enzyme-Linked Immunosorbent Assay (ELISA). Untreated cells were taken as a control. n = 3 independent experiments. Error bars represent the mean ± standard deviation. Two-tailed unpaired student’s t-test was performed to calculate statistical significance. (e). Image for the cytokine array shows IL-18 upregulation. A cytokine array was performed to measure the secretion of alisertib-induced senescent A549 cells after 72 hours of 2.5 µM salinomycin treatment. Untreated senescent cells were taken as a control. (f). Scatter plot showing log_10_(IC50) of salinomycin in alisertib-induced senescent A549 cells treated with different selected combinations of 10 µM Q-VD-OPh, Z-VAD-FMK, necrosulfonamide, MCC-950 for 7 days. Cells treated with salinomycin and without death pathway inhibitor treatment were taken as a control. (g). Western blot showing expression of DR5, JNK, p-JNK, cJUN and p-cJUN in alisertib-induced senescent A549 cells after 0, 24 and 48 hours of 2.5 µM salinomycin treatment with and without 5 mM ROS scavenger NAC. GAPDH served as a loading control. (h). Volcano plot showing differentially expressed JNK-cJUN cascade-induced proteins in alisertib-induced senescent A549 cells after 48 hours of 2.5 µM salinomycin treatment compared to untreated A549 cells. Protein abundance was measured using mass spectrometry. Thresholds for significance are 1.2 for abundance difference and 1.3 for -log_10_P. (i). To validate the knockdown efficiency, Real-time PCR analysis of cJUN was measured in A549 cells with two independent shRNAs against cJUN. pLKO empty vectors were taken as a control. n = 3 individual replicates. Error bars represent standard deviation. One-Way ANOVA was performed to calculate statistical significance. (j). Western blot showing DR5 expression in A549 cells with shRNA against cJUN after 48 hours of 2.5 µM salinomycin treatment. Untreated cells and pLKO empty vectors were taken as a control. Vinculin (VINC) served as a loading control.

Induction of senescence is associated with an increase in ROS levels (Fig. 2g). ROS are pleiotropic, affecting many different aspects of cell physiology.^25^ We treated senescent A549 cells with salinomycin with or without NAC and studied the effect of ROS scavenging on multiple pathways that we found to be activated by salinomycin. We found a significant reduction in DR5 and JNK-JUN signaling levels in the NAC-treated cells, highlighting the possibility that this pathway might be responsible for the DR5 activation (Fig. 3g). We also observed that salinomycin activates JNK-JUN cascade at transcriptomic and proteomic levels (Fig. 3h and AI appendix Fig. S3g). Therefore, we generated c-JUN knockdown A549 cells (Fig. 3i) and treated these cells with salinomycin. Our data indicate that the upregulation of DR5 was abrogated in the knockdown cells (Fig. 3j), indicating that salinomycin induces a strong upregulation of DR5 through ROS-mediated activation of the JNK-JUN signaling cascade.

### Salinomycin synergizes with DR5 agonist to facilitate tumor control through immune infiltration and activation

Our previous study revealed that the agonistic DR5 antibody conatumumab can serve as a senolytic agent, but the lack of activity and toxicity complicates clinical applications of DR5 agonistic antibodies.^16^ Building on our finding that salinomycin mediates DR5 induction, we investigated whether the synergistic effect of salinomycin and the DR5 agonist conatumumab could enhance elimination of senescent cells. We observed that the combination of salinomycin and conatumumab is strongly synergistic in senescent A549 cells (Fig. 4a). Essentially the same result was seen in additional cancer cell lines (AI Appendix Fig. S4a). We also measured the bliss synergy score of the combination to quantify the levels of synergy across a broader range of concentrations. With the salinomycin-conatumumab senolytic cocktail, we observed that the synergy bliss score is higher than 10 (indicating synergy) for many different senescent cancer cell models (Fig. 4a, b). We then tested the activity of the senolytic cocktail *in vivo* using subcutaneous injection of A549 cells in immunodeficient mice. In these experiments, we used alisertib as an inducer of senescence,^16,32^ and combinations of salinomycin and conatumumab as the senolytic cocktail. The concentration of alisertib for the *in vivo* experiments was chosen according to previous reports showing good senescence induction through p21 staining.^16^ Treatment with single-agent alisertib, salinomycin or conatumumab resulted in limited tumor growth inhibition. Treatment with the combination of two drugs resulted in some tumor control, but the strongest effect was seen with the three-way combination, without affecting bodyweight of the mice (Fig. 4c, AI appendix Fig. S4b). These findings underscore the efficacy of a combined approach utilizing salinomycin and conatumumab to target senescent cells, demonstrating a synergistic senolytic effect better to that of individual agents. Importantly, this cocktail does not show major toxicity levels when administered *in vivo*, suggesting its potential as a viable therapeutic strategy, but to evaluate the full toxicity profile of the cocktail further analyses are required.

**Figure 4.**
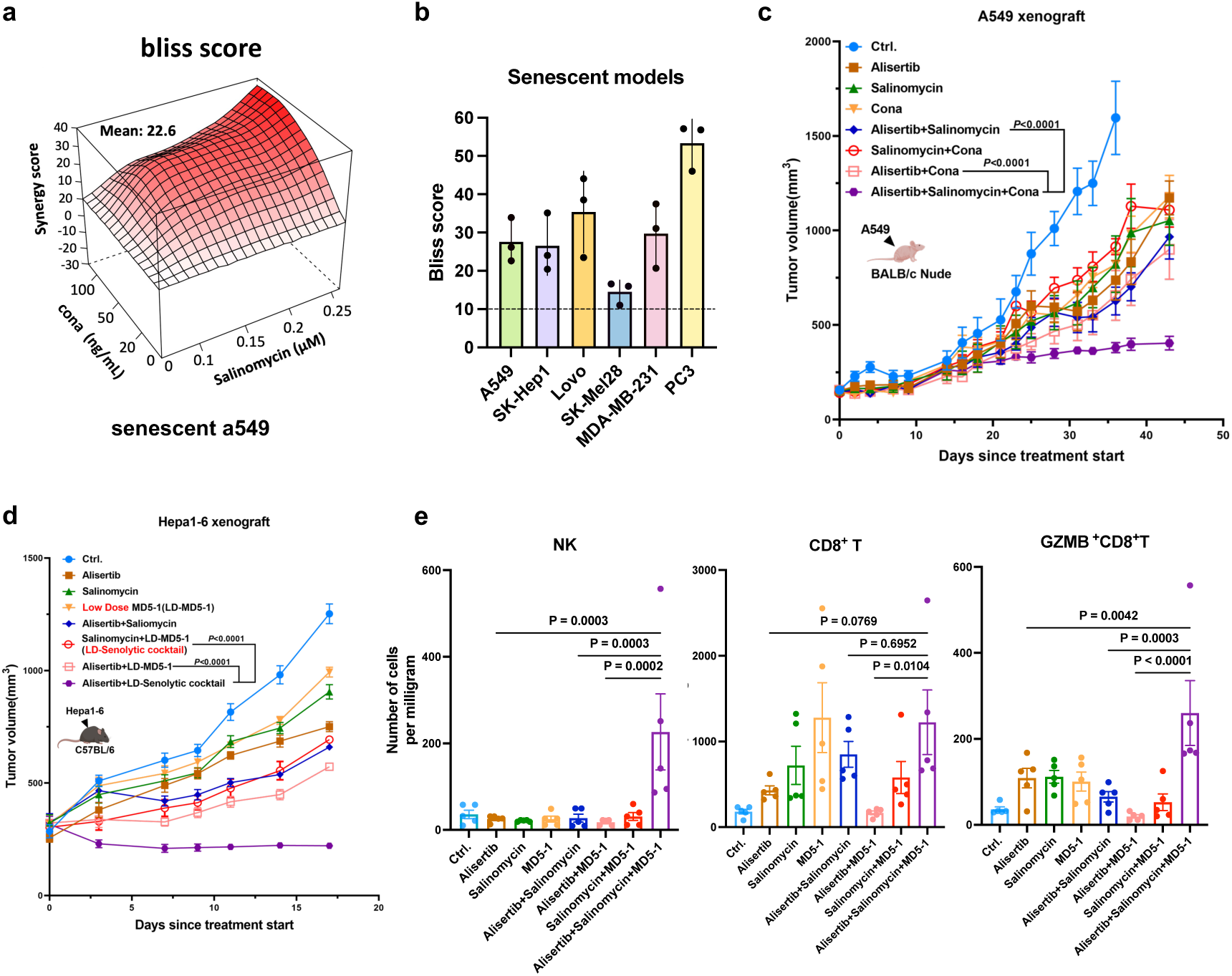
Salinomycin upregulates DR5 via ROS mediated JNK signaling activation and synergizes with DR5 agonist in vivo. (a). Bliss score 3D plots showing the synergy between DR5 agonist conatumumab and salinomycin in alisertib-induced senescent A549 (bliss score = 22.6). (b). Bar graph showing the bliss scores of the synergy between DR5 agonist conatumumab and salinomycin in alisertib-induced senescent A549, SK-Hep1, Lovo, SK-Mel-28, MDA-MB-231 and PC3 cells. A BLISS score above 10 is considered as synergistic. n = 3 independent experiments. Error bars represent the mean ± standard deviation. (c). Tumor growth of A549 lung cancer cells in the flank immunodeficient nude mice. When tumors reached approximately 100 mm^3^, animals were assigned to either vehicle control, 25 mg kg-1 alisertib, 1.5 mg kg-1 salinomycin, 5 µg per dose conatumumab (Cona), the two-way combinations of 1.5 mg kg-1 salinomycin plus 5 µg per dose conatumumab and 25 mg kg-1 alisertib plus 5 μg per dose conatumumab, or the three-way combination of 25 mg kg-1 alisertib plus 1.5 mg kg-1 salinomycin plus 2.5 µg per dose conatumumab. The body-weight of the mouse was monitored throughout the whole duration of the experiment (n = 7 mice per group). (d). Tumor growth of Hepa1-6 murine hepatoma cells in syngeneic immunocompetent C57BL/6 mice. When tumors reached approximately 250 mm^3^, animals were assigned to either vehicle control or 25 mg kg-1 alisertib as senescent inducer. The senolytic cocktail dosage was constant for the three experiments, equal to 1.5 mg kg-1 salinomycin, 0.3 µg per dose MD5-1, used as single or double agents and in a three-way combination with the different senescence inducers. N = 6 mice per group. (e). Quantitative estimate of various immune effector cells per milligram of tumor tissue in the tumors with different treatment groups, as determined by flow cytometry. n = 5 tumors per group.

Given the highly inflammatory phenotype of pyroptotic cell death, and the associated release of cytokines such as IL18 (Fig. 3e), we investigated the impact of our senolytic cocktail on immune cell responses to senescent cancer cells within the tumor microenvironment in an immune-competent context. We utilized a syngeneic model by inoculating aggressively growing Hepa1-6 murine liver cancer cells into C57BL/6 mice. Given the specificity of conatumumab for human DR5, we administrated the murine DR5 agonistic antibody MD5-1 in these experiments.^33^ As we suspected that the immune system would also play a role in tumor control, we applied a lower dose of the senolytic cocktail in this experiment. With a lower dose we would decrease the direct senolytic ability of the cocktail, while potentially introducing an immune mediated tumor control. Consistent with the previous findings, we observed a substantial tumor regression only for the combination of alisertib as a senescence inducer with the senolytic cocktail, without treatment associated body weight loss (Fig. 4d and AI Appendix Fig. S4c). Additionally, we observed superior tumor control as compared to the immune-deficient setting, which could not be readily explained by the senolytic activity of the compounds, as they were used in lower doses compared to the experiment in immune-deficient animals. The same tumor-controlling effect was observed using two chemotherapeutic drugs as senescence inducers: 5-FU and doxorubicin (AI Appendix Fig. S4d-g). The concentrations used for those two agents were selected according to previous reports showing proper senescence induction *in vivo* via p21 staining.^42,43,44^ Our data highlight that the senolytic cocktail has a widespread senolytic activity in different tumor models *in vivo*, which is also independent of the senescence inducer. Moreover, we collected the tumors and quantitative assessed the various immune cell populations using flow cytometry. Among the analysis, a notable infiltration of activated granzyme B+ and IFN-γ+ CD8 T cells, as well as NK cells were observed within the tumors following treatment with the senescence inducer in combination with the senolytic cocktail (Fig. 4e, AI appendix Fig. S4h). These results suggest that a better tumor control in immunocompetent animals, despite the use of a low-dose senolytic cocktail, are most likely the result of the activity of immune cells.

### Senolytic cocktail controls tumor growth via IL-18-mediated NK and CD8 T cell activation

To investigate the potential contributions of different immune cell populations to immune surveillance, we functionally inhibited two prominent groups of cytotoxic lymphocytes, NK1.1/CD161+ NK cells and CD8+ T cells, in an immunocompetent model using blocking antibodies. Our findings reveal that each antibody intervention attenuates anti-tumor efficacy of the triple drug combination to some extent. However, the concurrent blockade of both cytotoxic NK cells and CD8+ T cells result in the strongest abrogation of anti-tumor efficacy (Fig. 5a and AI appendix Fig. S5a). Finally, we investigated the interaction between the pyroptotic cytokine IL18 and our senolytic cocktail-mediated immune surveillance. IL18 blocking antibody was administered to tumor-bearing mice treated with the triple drug combination. We observed a dominant attenuation of tumor suppression and the blockade of activated T and NK cell infiltration (Fig. 5b, c and AI Appendix Fig. S5b). These findings suggest that IL18 plays an essential role in the senolytic-mediated immune surveillance in our model. These results collectively suggest that both cytotoxic NK cells and CD8+ T cells attracted by the pyroptotic cell death cytokine IL18, play dominant roles in orchestrating senolysis-mediated by the drug cocktail used here. Together, we advocate a therapeutic strategy to combine pro-senescence therapy with our senolytic cocktail, which can elicit a multifaceted response, instigating diverse cell death mechanisms and bolstering immune surveillance (Fig. 5d).

**Figure 5.**
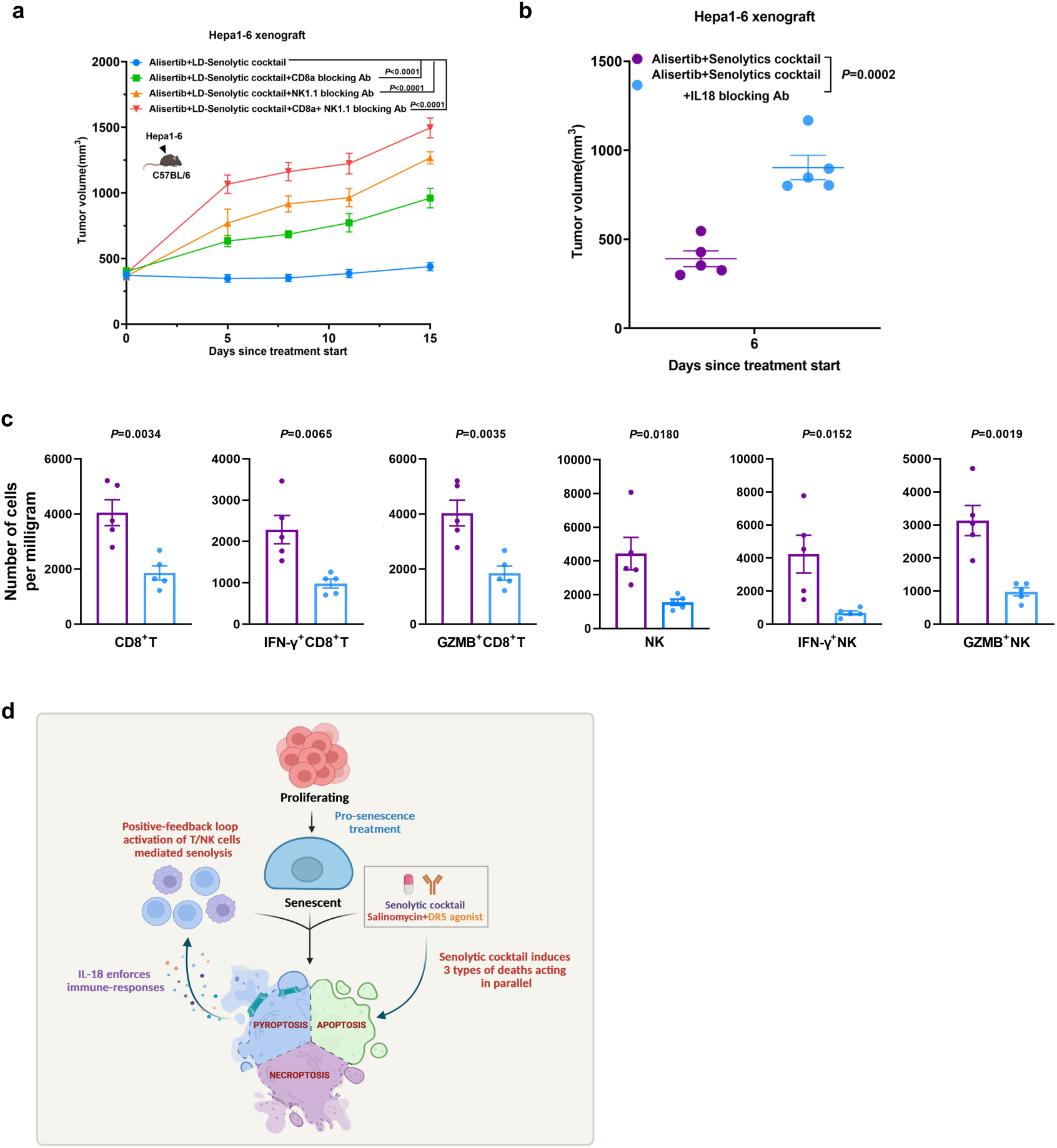
NK and CD8+ T cells modulate the tumor control thanks to the IL18 mediated immune recruitment and activation in vivo. (a). Tumor growth of Hepa1-6 murine hepatoma cells in syngeneic immunocompetent C57BL/6 mice. When tumors reached approximately 300 mm^3^, animals were intraperitoneally (i.p.) injected with 200 µg anti-CD8a, 200 µg anti-NK1.1 or the combination, one day before three-way combination of 25 mg kg-1 alisertib plus 1.5 mg kg−1 salinomycin plus 0.3 µg per dose MD5-1, and then the treatment was continued on days 2, 5, and 8 with lower dose of 100 µg anti-CD8a and anti-NK1.1 antibodies (n = 5 mice per group). (b). Tumor growth of Hepa1-6 murine hepatoma cells in syngeneic immunocompetent C57BL/6 mice. When tumors reached approximately 200 mm^3^, animals were intraperitoneally (i.p.) injected with 50 µg anti-IL 18, three-way combination of 25 mg kg-1 alisertib plus 1.5 mg kg−1 salinomycin plus 0.3 µg per dose MD5-1, and then the treatment was continued on days 3 with 50µg anti-IL 18 (n = 5 mice per group). (c). Quantitative estimate of various immune effector cells per milligram of tumor tissue in three-way combination and three-way combination+IL 18 blocking antibody treated Hepa1-6 tumors, as determined by flow cytometry. n = 5 tumors per group. NK, natural killer cell. (d). The model involving pro-senescence therapy and a senolytic cocktail elicits a multifaceted response, instigating diverse cell death mechanisms and bolstering immune surveillance.

## Discussion

Many cancer drugs induce senescence in tumor cells, leading to chronic inflammation. However, the lack of effective senolytic agents limits the possibilities of eliminating them. We show here that the antibiotic salinomycin, combined with DR5 agonistic antibody, exhibits potent senolytic activity. A genome-wide CRISPR screen identified SLC25A23 depletion as a vulnerability in senescent cancer cells. The absence of direct inhibitors for this protein led us to perform a drug screen with drugs that target SLC25A23 functions and/or its known interactors, such as the mitochondrial calcium uniporter MCU. This resulted in the identification of salinomycin, an antibiotic that triggers multiple stress responses, leading to cell death pathways including apoptosis, necroptosis, and pyroptosis. The occurrence of three death mechanisms acting in parallel is unusual, and usually killing occurs through one more dominant cell death pathway. However, we demonstrate here that all three pathways play a part in the senolytic effect of salinomycin. We show that single death inhibitors cannot reduce killing, but only by using inhibitors of all three pathways simultaneously can we fully prevent salinomycin senolytic effect, highlighting the independent occurrence of these pathways. It is important to note that increasing evidence is also showing that the different death pathways are highly interconnected, mostly due to the capability of the caspases to cross-activate each other, but the choice of one death type over the other and the kinetics by which this cross-activation occurs is still understood poorly.

Cancer patients that are exposed to the detrimental chronic effects of SASP produced by senescent cells generally have worse prognosis. Clinical data have shown that the presence of senescence correlates with lower overall survival^34,35^. This highlights the clear clinical need to develop therapies that can induce the death of senescent cells. Previously described senolytic therapies, namely DR5 targeting antibodies and ABT-263, generally share the same restrictions, as they have insufficient activity across different tumor types, and exhibit notable toxicities in clinical settings. Here, we propose a cocktail of drugs that can efficiently eradicate a variety of senescent tumor cells, regardless of the senescence inducers in vitro and in vivo. Of potential clinical relevance is our finding that the anti-cancer effects of the chemotherapeutic agents doxorubicin and 5-FU can be potentiated by combining with the salinomycin+DR5 agonist senolytic cocktail, without strong body-weight loss in mouse. Chemotherapeutic agents usually induce apoptosis in actively proliferating cells, leaving mostly unaffected cells that withdraw from the cell cycle, such as the senescent cells. The use of combinations of chemotherapy and senolytics can be particularly favorable in the clinic to kill senescent cells.

The immune cell effects triggered by the senolytic cocktail allow the use of lower concentrations of senolytic drugs in immunocompetent animals, thereby reducing potential toxicities. The concentration of salinomycin required to achieve strong synergy with conatumumab is significantly lower than previously reported.^45^ Our findings underscore the importance of concurrently targeting the multiple vulnerabilities of senescent cells to increase cell death while reducing the emergence of resistant cells. Notably, two of the death mechanisms occurring in our model, namely necroptosis and pyroptosis, are recognized as immunogenic programmed types of death. We demonstrate that salinomycin-treated senescent cells increase the secretion of the immunostimulatory cytokines IL1β and IL18, derived from the induction of pyroptosis. In a previous study, the senolytic effect of activating DR5 through the agonistic antibody conatumumab was shown.^16^ Here, we advance this finding by showing that the combination of salinomycin with conatumumab not only exhibits stronger senolytic activity in vitro and in vivo, but also induces higher NK and CD8+ T cell infiltration and activation compared to single agents when combined with senescence inducers. We also provide proof that by further reducing the concentration of the senolytic cocktail in vivo, we still achieve strong tumor control, primarily mediated by the killing activity of the NK and CD8+ T cells. This effect is mostly due to the release of the pyroptotic cytokine IL18. However, it remains unclear why the potentiation of apoptosis through DR5 activation would increase the pyroptosis-induced release of IL18 by salinomycin and immune cell infiltration and activation. One possibility could be that during the late stages of apoptosis, stronger cell lysis could increase cytokine release. This immune effect is superimposed on the well-established inflammatory effects induced by the SASP of senescent cells. Our senolytic cocktail could offer therapeutic opportunities by inducing cancer cell death and reducing detrimental effects of the chronic SASP by senescent cells eradication. Effector cytotoxic T cells and NK cells mostly induce cancer cell death through extrinsic apoptosis pathway activation, which is strongly enhanced in our setting by the salinomycin-induced DR5 upregulation. Finally, several DR5-agonistic antibodies have exhibited notable toxicity and limited efficacy in clinical settings. Combining lower doses of DR5 agonistic antibodies with salinomycin offers a potential therapeutic approach to reduce toxicity while maintaining efficacy in the clinic.

## Materials and Methods

### Cell lines

All cell lines were obtained from ATCC. Cell lines A549 (CCL-185), SK-Mel28 (HTB-72), MB-MDA-231 (HTB-26), Lovo (CCL-229) and PC3 (CRL-1435) were cultured in a RPMI-based medium supplemented with 10% FBS, 1% penicillin/streptomycin and 2 mM L-glutamine. Cell lines SK-Hep1 (HTB-52), T47D (HTB-133) and MCF-7 (HTB-22) were cultured in a DMEM-based medium supplemented with 10% FBS, 1% penicillin/streptomycin and 2 mM L-glutamine. Cell line HEK-293 (CRL-1573) was cultured in a DMEM-based medium supplemented with 10% FBS and 1% penicillin/streptomycin. Monoclonal A549-iCas9 were generated from A549 by infecting the cells with the edit-R Inducible Lentiviral Cas9 vector. All the cell lines have been validated by STR profiling and regularly tested for Mycoplasma contamination with PCR-based assay.

### Compounds and antibodies

Alisertib (#201931), barasertib (#200420), etoposide (#100330), PF-06873600 (#206997), Q-VD-OPh, (#526845), Z-VAD-FMK (#471029), necrosulfonamide (#525389), MCC950 (#522637) and 3-methyladenine (#406971) were purchased from MedKoo Biosciences Inc. CFI-400945 (#S7552) was purchased from Selleck Chemicals. Salinomycin (#HY-15597) and Nigericin (#HY-127019) were purchased from Medchem Express. The InVivoMAb anti-mouse DR5 (MD5-1) antibody (#BE0161) and InVivoMAb polyclonal Armenian hamster IgG (#BE0091) was used as the isotype control of MD5-1 and was purchased from BioXcell. Anti-vinculin (hVIN-1, #V9131) was purchased from Sigma-Aldrich. Anti-DR5 (D4E9, #8074), anti-GAPDH (D16H11, #5174), anti-c-Jun (L70B11, #2315), anti-phospho-c-Jun (Ser73) (D47G9, #3270), anti-cleaved gasdermin D (Asp275) (E7H9G, #36425), anti-cleaved caspase 3 (Asp175) (polyclonal, #9661), anti-MLKL (D2I6N, #14993), anti-phospho-MLKL (Ser358) (D6H3V, #91689), anti-LC3B (#2775) and anti-P21 (polyclonal, #2947) were purchased from Cell Signaling Technology. Anti-caspase 1 (14G468, #sc-56036), anti-JNK (sc-571), anti-phospho-JNK (Thr 183 / Tyr 185) (polyclonal, #sc-12882) and anti-HSP90 (polyclonal, #sc-7947) were purchased from Santa Cruz Biotechnology. Anti-gasdermin D (EPR19828, #ab209845) was purchased from Abcam. Secondary antibodies Goat Anti-Mouse IgG (#1706516) and Goat Anti-Rabbit IgG (#1706515) were purchased from Bio-Rad. Swine Anti-Goat IgG (#MBS674933) was purchased from MyBioSource. DR5 agonistic antibody conatumumab was purchased from UMAB. Murine specific anti-DR5 agonistic antibody MD5-1 (#MD5-1), anti-CD8a (#2.43) and anti-NK1.1 (#PK-136) were purchased from BioXcell.

### Plasmids and primers

All the primers and individual shRNA vectors were collected from the TRC library. gRNA cloning vector pLentiGuide-puro (#52963)^36^ and Human CRISPR Knockout Pooled Library (Brunello) (#73178)^37^ were purchased from Addgene. The edit-R Inducible Lentiviral Cas9 vector (#NC1606271) was purchased from Dharmacon.

### iCas9-associated senolytic CRISPR screen

A549 were infected with a doxycycline-inducible Cas9 (iCas9) lentiviral vector to generate an inducible Cas9 cell model (A549-iCas9). Next, the Brunello lentiviral whole genome-wide gRNA collection virus was introduced to the A549-iCas9 cells. The infected cells were firstly treated with 0.5 µM alisertib for 7 days to induce senescence. Afterward, these cells were suspended from alisertib and then switched to 1 µg/ml doxycycline treatment for 10 days. Non-senescent cells were included as the control arm to filter out the straight lethal genes and treated with doxycycline. Changes in library representation after 10 days of doxycycline treatment were determined by Illumina deep-sequencing. gRNAs prioritized for further analysis were selected by the fold depletion of abundance in the S2 and S1 samples compared with that in P2 and P1 samples, using methods as described previously.^16,38^

### Drug screen

To identify the most promising senolytic drug, a drug screen was performed using parental and senescent A549 cells. The drug screen included OXPHOS inhibitors FCCP (#522447), Pyrvinum Pamoate (#HY-A0293), Antimycin A (#527667), Atovaquone (#317258), Fenofibrate (#317866), Devimistat (#200823), Mitoxantrone (#HY-13502), Celastrol (#200694), Elesclomol (#201210), VLX600 (#408097), Bay-872243 (#205901), Metformin (#575896), Canagliflozin (#314203) and Rotenone (#581167), the mithcondrial function inhibitors Tigecyclin (#315297), Doxycycline (#592623), Mupirocin (#318292), Docetaxel (#100270), Cisplatin (#100160) and Actinonin (#406221), the ER stress modulators Thapsigargin (#561537), 2-ABP (#1224), Carbachol (#PHR1511) and Chloroquine (#598387), and the calcium regulators Ru360 (#557440), Benzethonium chloride (#531006), Amorolfine (#461377), Ionomycin (#540223), Salinomycin (#406545), BAPTA-AM (#563565), Amlodipine (#317184), Cinacalcet (#315239), Furosemide (#317960), BAYK 8644 (#531565), Diltiazem (#317617), Nifedipine (#318364), Berbamine hydrochloride (#471219) and Verapamil (#592564). ABT-263 (#201970) was included as a positive control. Pyrvinium Pamoate and Mitoxantre were purchased from MedChemExpress. 2-APB was purchased from bio techne Tocris. Carbachol and Ru360 were purchased from Sigma-Aldrich. All other drugs were purchased from MedKoo Biosciences Inc. Cells were treated once with a range of 0.01 µM to 20 µM of the drugs, and cell viability was assessed by CellTiter-Blue assay five days after treatment. The senolytic index of the drugs was calculated by dividing the IC50 of the drugs in parental A549 cells by the IC50 in senescent A549 cells.

### Drug treatments

Senescence was induced in MDA-MB-231, SK-Hep1 and A549 cells by treating the cells with 0.5 µM alisertib, 1 µM barasertib, 2 µM etoposide, 100 nM CFI-400945 or 0.5 µM PF-06873600 for 7 days. In case of additional treatment, cells were washed with PBS, trypsinized and reseeded. The concentrations used for salinomycin, conatumumab, nigericin, antimycin A, and rotenone treatment in vitro are indicated in the Figs. For NAC treatment, 5 mM was added 24 hours prior to the start of salinomycin treatment and cells were retreated with NAC every other day. For death pathway inhibitors Q-VD-OPh, Z-VAD-FMK, necrosulfonamide, MCC-950 and 3-methyladenine (3-MA) 10 µM was added 4 hours prior to salinomycin treatment. To validate Salinomycin as senolytic agent we seeded the cells at day zero and started treatment at day one and every 3 days for a duration of one week experiment (two treatments total). Media was changed every time the drug was refreshed. For proteomics and bulk RNA seq experiments cells were seeded in appropriate number and treated with salinomycin for 48 hours. Cells were then collected for protein and RNA extraction and processed as indicated above in the dedicated sections of the methods. For the senolytic cocktail experiments cells were seeded in 384 well plate and treated with a single treatment of both salinomycin and conatumumab. Media was refreshed every 3 days and viability was measured at day 5 using cell titer blue. Dose-response curves and IC50 values were calculated using GraphPad Prism 10. The concentrations used for in vivo experiments are indicated in the figures.

### Staining for senescence-associated β-galactosidase activity

β-galactosidase activity in cells was detected using a Histochemical Staining kit (CS0030-1KT) from Sigma-Aldrich. β-galactosidase detection was carried out according to the manufacturer’s instructions. SA-β-gal staining-positive cells were quantified based on three independent images.

### Cell viability measurements

Cell viability was assessed by either CellTiter-Blue assay or Colony Formation Assay. 1k-20k cells were seeded in each experiment according to cell growth rate and well surface area. For CellTiter-Blue assay the cells were treated with 20x cell titer blue (Thermofisher #R12204) and cell viability was measured with EnVision multi-label plate reader (PerkinElmer). For Colony Formation Assay the cells were fixed with 4% paraformaldehyde (Merck Millipore #104002) diluted in PBS and stained with 2% crystal violet (Sigma-Aldrich #HT90132) diluted in water. Pictures were taken with Epson perfection V750 scanner, and the data was quantified using Fiji.

### Synergy plots

Senescent A549, SK-Hep1, Lovo, SK-Mel28, MDA-MB-231, and PC3 cells were treated in a dose-response matrix with an increasing concentration of salinomycin on one axis and an increasing concentration of DR5 agonist conatumumab on the other axis for 7 days. Cell viability was assessed with the CellTiter-Blue assay. Synergy plots were generated, and bliss scores were calculated using SynergyFinder (https://www.synergyfinderplus.org/).

### Lentiviral transduction

A third-generation lentivirus packaging system consisting of pCMV-VSV-G (Addgene 8454), pRSV-Rev (Addgene 12253) and pMDLg/ pRRE (Addgene 12251) was used to create virus particles of the modified reporter plasmids. A transient transfection was performed in 293T cells and lentiviral supernatants were produced. Destination cells were infected with lentiviral supernatants, using 8 μg ml−1 polybrene and low virus titer. After 48 h of incubation, the supernatant was replaced by a medium containing 10 μg ml−1 blasticidin or 2 μg ml−1 puromycin. After 48 h, the selection of viral transduced cell lines was completed.

### Protein lysates generation and Western blot

After treatment, cells were washed with PBS and lysed with RIPA buffer supplemented with protease inhibitors (cOmplete, Roche) and phosphatase inhibitor cocktails II and III (#P5726 and #P0044, Sigma-Aldrich). All lysates were freshly prepared and processed with Novex NuPAGE Gel Electrophoresis Systems (Invitrogen). The detection was performed after 48- or 72-hours drug treatment. Following the specified culture duration and drug treatments, cells underwent cold phosphate-buffered saline (PBS) washes and lysis in RIPA buffer (25 mM Tris-HCl, pH 7.6, 150 mM NaCl, 1% NP-40, 1% sodium deoxycholate, and 0.1% SDS) supplemented with complete protease inhibitor (Roche) and phosphatase inhibitor cocktails II and III (Sigma). The lysates were then clarified by centrifugation at 15,000 × g for 30 minutes at 4 °C and quantified using a Bicinchoninic Acid assay (Pierce BCA, Thermo Fisher Scientific), following the manufacturer’s protocol. Protein samples were denatured with DTT and heated at 100 °C for 5 minutes. Approximately 20 µg of protein samples were loaded and subjected to SDS-PAGE for about 1 hour at 150 V, followed by transfer onto a nitrocellulose membrane at 350 mA for 120 minutes. Subsequently, the membranes were immersed in 5% BSA in TBS with 0.1% Tween-20 (TBS-T) for 1 hour for blocking. Following this, the membranes were incubated overnight at 4 °C with the designated primary antibodies in 5% BSA (Bovine Serum Albumin) in TBS-T. After three 5-minute washes with TBS-T, the membranes were incubated at room temperature for 1 hour with anti-rabbit or anti-mouse secondary antibodies (HRP conjugated) in 5% BSA in TBS-T. Following another three 5-minute washes in TBS-T, a chemiluminescence substrate (ECL, Bio-Rad) was applied to the membranes, and the signal was captured using the ChemiDoc-Touch system (Bio-Rad).

### qRT-PCR

Total RNA was isolated from cells utilizing either TRIzol reagent from Invitrogen or Quick-RNA™ MiniPrep (#R1055) from Zymo Research. Subsequently, cDNA synthesis was carried out employing the Maxima Universal First Strand cDNA Synthesis Kit (#K1661) from Thermo Scientific. For quantitative PCR (qPCR), FastStart Universal SYBR Green Master (Rox) from Roche was utilized. All procedures strictly adhered to the respective manufacturer’s protocols. Each reaction was run in duplicates or triplicates, and relative mRNA levels were determined using the Standard Curve Method. Primers sequences are provided below.

GAPDH Forward: AAGGTGAAGGTCGGAGTCAA;

GAPDH Reverse: AATGAAGGGGTCATTGATGG;

SLC25A23 Forward: AGGAATACCTGGCATTCCAG;

SLC25A23 Reverse: CCAAGGAGAGCACCTTGAAA;

cJUN forwards: TCCAAGTGCCGAAAAAGGAAG;

cJUN reverse: CGAGTTCTGAGCTTTCAAGGT;

### Bulk RNA sequencing

Sequencing data was demultiplexed into FastQ files using BCLConvert version 4.1.5 (Illumina). The paired-end reads were trimmed for adapter sequences using SeqPurge version 2019_09^39^ and aligned to the human reference genome using HiSat2version 2.1.0 with the pre-built grch38_snp_tran reference.^40^ After alignment, UMI aware duplicates were marked using Rumidup. The de-duplicated gene counts were determined by GenSum using Ensembl release 102 transcript GTF as a reference.^41^

### Seahorse Assay

Seahorse assay was performed using Agilent Seahorse XFe24 Analyzer according to the manufacturer’s instructions (Agilent). Concentrations of 10 µM FCCP (#522447), 10 µM Oligomycin A (#329825), 10 µM Antimycin (#527667) and 10 µM Rotenone (#581167) were used. All four compounds were purchased from MedKoo Biosciences Inc.

### ROS staining

ROS measurement was performed using the CellROX™ Green Flow Cytometry Assay Kit according to the manufacturer instruction. Cells were pretreated for 2 hrs with NAC to deplete ROS levels as negative control and with TBHP to induce high levels of ROS as positive control. Results were calculated by normalizing the ROS+ % to the positive and negative controls. The fluorescence of the CellROX™ Green dye was measured through flow cytometry using 488nm as excitation wavelength and the 530/30 emission filter.

### Calcium staining

Cells were stained with 1µM Rhod-2-AM calcium specific dye (Invitrogen, #R1244) for 1 hr at 37 C 5% CO_2_. Cells were then washed, trypsinized and resuspended in FACS-buffer supplemented with live dead staining DAPI prior to flow cytometry analysis. Fluorescence was measured using the 561nm excitation wavelength with 586/15 emission filter.

### Flow cytometry

Cells were either stained with CellROX Green for ROS Visualization or with Rhod-2 for calcium 17isualization. Cells were washed with PBS, trypsinized and resuspended in a phenol-red negative RPMI-based medium (ThermoFisher #11835030). Flow cytometry was performed with BD LSRFortessa. Results were analyzed by FlowJo 10.8.2.

### IL1β release

After senescence induction, A549 cells were treated with 2.5 µM salinomycin for 72 hours. The supernatant was collected. ELISA was performed using IL-1 beta Human ELISA Kit (Invitrogen #KHC0011) according to the manufacturer’s instructions.

### Cytokine array

After senescence induction, A549 cells were treated with 2.5 µM salinomycin for 72 hours. The supernatant was collected. A cytokine array was performed using the Proteome Profiler Human Cytokine Array Kit (Biotechne R&D systems #ARY005B) according to the manufacturer’s instructions.

### Proteomics

Cell pellets were lysed and sonicated in 1x S-Trap lysis buffer (5% SDS in 50mM TEAB, pH 8.5), after which aliquots comprising 30 µg of protein were reduced with 20 mM DTT (20 min. at 55°C) and alkylated with 40mM iodoacetamide (30 min. at RT in the dark). Proteins were digested at 37°C overnight with trypsin (Sigma-Aldrich; enzyme/substrate ratio 1:10) on S-Trap Micro spin columns according to the manufacturer’s instructions (ProtiFi, NY, USA). Peptides were eluted, vacuum dried and stored at -80°C until LC-MS/MS analysis. Nano-LC-MS/MS was performed on an Orbitrap Exploris 480 mass spectrometer (Thermo Scientific) connected to a Proxeon nLC1200 system. Peptides were directly loaded onto the analytical column (ReproSil-Pur 120 C18-AQ, 1.9μm, 75 μm × 500 mm, packed in-house) and eluted in a 90-minutes gradient containing a linear increase from 5% to 30% solvent B (solvent A was 0.1% formic acid/water and solvent B was 0.1% formic acid/80% acetonitrile). The Exploris 480 was run in data-independent acquisition (DIA) mode, with full MS resolution set to 120,000 at m/z 200, MS1 mass range was set from 350-1400, normalized AGC target was 300% and maximum IT was 45ms. DIA was performed on precursors from 400-1000 in 48 windows of 13.5 Da with an overlap of 1 Da. Resolution was set to 30,000 and normalized CE was 27.

RAW files were analyzed with DIA-NN (version 1.8) using standard settings. Fragment spectra were searched against the Swissprot human database (version 2022_08; 20,398 entries) by selecting ‘FASTA digest for library-free search’; Trypsin/P was specified as protease specificity allowing a maximum of 1 miscleavage; N-terminal excision (M) and carbamidomethylation (C) were selected as fixed modifications and match between runs was applied. Protein group abundances were extracted from the DIA-NN result files, imported into Perseus (2.0.10.0) and Log2-transformed. For group-wise comparisons and volcano plots, values were filtered for presence in at least 2 out of 3 replicates in at least one sample group. Missing values were replaced by an imputation-based normal distribution using a width of 0.3 and a downshift of 2.4. Differentially expressed proteins were determined using a student’s t-test p<0.05 and Log2 abundance ratio < -1 ^ >1 as cutoff values.

### Scanning Electron Microscopy

Fixed cells on glass coverslips were washed with water and dehydrated in an ethanol series: 30%, 50%, 70%, 80%, 90%, 96%, 100% each for 10 min, and subsequently in and acetone series: 30%, 50%, 100% each for 10 min. Samples were critical point-dried using a Leica EM CPD300 and mounted on aluminum stubs using carbon adhesive stickers. Subsequently a 6 nm Platinum / Palladium coating was applied using a Leica ACE 600 sputter coater. Samples were imaged using a Zeiss SEM, Sigma 300 Field Emission, Gemini with at 3 kV and Secondary Electron-2 detector.

### Cell line-derived xenografts

All animals were manipulated according to protocols approved by Sun Yat-Sen University. The maximal tumor size/burden was 2,000 mm^3^. All in vivo experiments did not exceed the maximal tumor size. Female 5–6-week-old BALB/c nude mice were used in the studies. As shown in Fig. 5a, A549 cells (1 × 10^7^ cells per mouse) were injected subcutaneously into the flanks of nude mice (7 mice per group). After tumor establishment, mice were randomly assigned to 5 days per week treatment (Drug-off in the weekend) and given vehicle, conatumumab (intraperitoneal injection), alisertib (oral gavage), salinomycin (oral gavage) or combinations. As shown in Fig. 5b, Hepa1-6 murine hepatoma cells (5 × 10^6^ cells per mouse) were injected subcutaneously into the flanks of in syngeneic immunocompetent C57BL/6 mice (6 mice per group). After tumor establishment, mice were randomly assigned to 5 days per week treatment (Drug-off in the weekend) and given vehicle, alisertib, salinomycin or related combinations. In the treatment arms involved murine DR5 agonist MD5-1, MD5-1 were intraperitoneally injected 5 days per week treatment (Drug-off in the weekend). As shown in Fig. 5c, Hepa1-6 cells (5 × 10^6^ cells per mouse) were injected subcutaneously into the flanks of syngeneic immunocompetent C57BL/6 mice (5 mice per group). After tumor establishment, mice were randomly assigned to 5 days per week treatment (Drug-off in the weekend) and given vehicle, 5-fluorouracil, salinomycin or related combinations. In the treatment arms involved mouse DR5 agonist MD5-1, MD5-1 were intraperitoneally injected 5 days per week treatment (Drug-off in the weekend). As shown in Fig. 5d, Hepa1-6 cells (5 × 10^6^ cells per mouse) were injected subcutaneously into the flanks of syngeneic immunocompetent C57BL/6 mice (5 mice per group). After tumor establishment, mice were randomly assigned to 5 days per week treatment (Drug-off in the weekend) and given vehicle, doxorubicin, salinomycin or related combinations. In the treatment arms involved murine DR5 agonist MD5-1, MD5-1 were intraperitoneal injected 5 days per week treatment (Drug-off in the weekend). For Fig. 5a, tumor growth of Hepa1-6 cells were injected in completely immune-incompetent NCG mice. When tumors reached approximately 250 mm^3^, animals were assigned to either vehicle control, or the three-way combination of 25 mg kg-1 alisertib plus 1.5 mg kg−1 salinomycin plus 0.3 µg per dose MD5-1 (5 mice per group). For blocking NK and T cell in vivo, as shown in Fig. 5f, Hepa1-6 cells (5 × 10^6^ cells per mouse) were injected subcutaneously into the flanks of in syngeneic immunocompetent C57BL/6 mice (5 mice per group). After tumor establishment, mice were randomly assigned to 5 days per week three-way combination treatment (Drug-off in the weekend). In the treatment arms involved murine DR5 agonist MD5-1, MD5-1 were intraperitoneal injected 5 days per week treatment (Drug-off in the weekend). Tumor volume based on calliper measurements was calculated by the modified ellipsoidal formula of tumor volume = ½ length × width^2^. Data are presented as mean ± s.e.m. Mice that showed therapy-induced adverse events in the early phases of treatment, precluding conclusions on the treatment efficacy, were excluded from longitudinal analyses. Exclusion criteria were predetermined before the experiments.

### IL18 and T, NK cell correlation

Transcriptomic data and clinical information from TCGA LIHC cohort were downloaded from the UCSC Xena data portal (https://xenabrowser.net). Microenvironment Cell Populations-counter (MCP-counter) was used for estimating the absolute abundance of CD8 T cells and NK cells in each TCGA tissue sample.

## Supporting information

Supplemental figures Casagrande

## Statistics and reproducibility

Statistical analysis was performed using Graph Pad Prism 10. Statistical analysis was carried out on the data from independent experiments. The error bars in the Figures represent s. d.; the n value is specified in the legends.

## Resource availability

Requests for further information and reagents may be addressed to the corresponding author: Rene Bernards or Liqin Wang.

## Data and code availability

All omics data will be uploaded to GEO once manuscript being conditional accepted, and will be listed in the key re-sources table.

## Acknowledgments

We thank members of the Wang, Bernards and Beijersbergen laboratories for helpful discussion and insightful feedback. We thank the genomics core facility for the RNAseq experiments and sample processing. This work was supported by the European Research Council as ERC-787925 (R.B.), 19-051-ASP from the Mark Foundation (R.B.), ASP-II grant-Bernards 2023 (R.B.), KWF-12539 from the Dutch Cancer Society (R.B.), LSH-TKI-LSHM20083 from Health Holland (R.B.); the X-omics Initiative (Project 184.034.019), part of the NWO National Roadmap for Large-Scale Research Infrastructures (O.B.B.); W2432050; 82372695 from the National Natural Science Foundation of China (L.W.) and 2024A04J6484 from the Guangdong Basic and Applied Basic Research Foundation (L.W.).

## References

1. Hayflick, L., and Moorhead, P.S. (1961). The serial cultivation of human diploid cell strains. Exp. Cell Res. 25, 585–621. 10.1016/0014-4827(61)90192-6.

2. Wang, L., Lankhorst, L., and Bernards, R. (2022). Exploiting senescence for the treatment of cancer. Nat. Rev. Cancer 22, 340–355. 10.1038/s41568-022-00450-9.

3. Coppé, J.P., Desprez, P.Y., Krtolica, A., and Campisi, J. (2010). The senescence-associated secretory phenotype: The dark side of tumor suppression. Annu. Rev. Pathol. Mech. Dis. 5, 99–118. 10.1146/annurev-pathol-121808-102144.

4. Coppé, J.P., Patil, C.K., Rodier, F., Sun, Y., Muñoz, D.P., Goldstein, J., Nelson, P.S., Desprez, P.Y., and Campisi, J. (2008). Senescence-associated secretory phenotypes reveal cell-nonautonomous functions of oncogenic RAS and the p53 tumor suppressor. PLoS Biol. 6. 10.1371/journal.pbio.0060301.

5. Prata, L.G.P.L., Ovsyannikova, I.G., Tchkonia, T., and Kirkland, J.L. (2018). Senescent cell clearance by the immune system: Emerging therapeutic opportunities, 10.1016/j.smim.2019.04.003 10.1016/j.smim.2019.04.003.

6. Ruscetti, M., Morris, J.P., Mezzadra, R., Russell, J., Leibold, J., Romesser, P.B., Simon, J., Kulick, A., Ho, Y. jui, Fennell, M., et al. (2020). Senescence-Induced Vascular Remodeling Creates Therapeutic Vulnerabilities in Pancreas Cancer. Cell 181, 424–441.e21. 10.1016/j.cell.2020.03.008.

7. Batlle, E., and Massagué, J. (2019). Transforming Growth Factor-β Signaling in Immunity and Cancer. Immunity 50, 924–940. 10.1016/j.immuni.2019.03.024.

8. Faget, D. V., Ren, Q., and Stewart, S.A. (2019). Unmasking senescence: context-dependent effects of SASP in cancer, 10.1038/s41568-019-0156-2 10.1038/s41568-019-0156-2.

9. Niedernhofer, L.J., and Robbins, P.D. (2018). Senotherapeutics for healthy ageing, 10.1038/nrd.2018.44 10.1038/nrd.2018.44.

10. Schmitt, C.A., Wang, B., and Demaria, M. (2022). Senescence and cancer — role and therapeutic opportunities. Nat. Rev. Clin. Oncol. 19, 619–636. 10.1038/s41571-022-00668-4.

11. Xu, M., Pirtskhalava, T., Farr, J.N., Weigand, B.M., Palmer, A.K., Weivoda, M.M., Inman, C.L., Ogrodnik, M.B., Hachfeld, C.M., Fraser, D.G., et al. (2018). Senolytics improve physical function and increase lifespan in old age. Nat. Med. 24, 1246–1256. 10.1038/s41591-018-0092-9.

12. Baker, D.J., Childs, B.G., Durik, M., Wijers, M.E., Sieben, C.J., Zhong, J., A. Saltness, R., Jeganathan, K.B., Verzosa, G.C., Pezeshki, A., et al. (2016). Naturally occurring p16 Ink4a-positive cells shorten healthy lifespan. Nature 530, 184–189. 10.1038/nature16932.

13. Schmitt, C.A., Fridman, J.S., Yang, M., Lee, S., Baranov, E., Hoffman, R.M., and Lowe, S.W. (2002). A senescence program controlled by p53 and p16INK4a contributes to the outcome of cancer therapy. Cell 109, 335–346. 10.1016/S0092-8674(02)00734-1.

14. Carpenter, V.J., Saleh, T., and Gewirtz, D.A. (2021). Senolytics for cancer therapy: Is all that glitters really gold?, 10.3390/cancers13040723 10.3390/cancers13040723.

15. Paez-Ribes, M., González-Gualda, E., Doherty, G.J., and Muñoz-Espín, D. (2019). Targeting senescent cells in translational medicine. EMBO Mol. Med. 11. 10.15252/emmm.201810234.

16. Wang, L., Jin, H., Jochems, F., Wang, S., Lieftink, C., Martinez, I.M., De Conti, G., Edwards, F., de Oliveira, R.L., Schepers, A., et al. (2022). cFLIP suppression and DR5 activation sensitize senescent cancer cells to senolysis. Nat. cancer 3, 1284–1299. 10.1038/s43018-022-00462-2.

17. Görlach, A., Bertram, K., Hudecova, S., and Krizanova, O. (2015). Calcium and ROS: A mutual interplay. Redox Biol. 6, 260–271. 10.1016/j.redox.2015.08.010.

18. Bertero, E., and Maack, C. (2018). Calcium signaling and reactive oxygen species in Mitochondria. Circ. Res. 122, 1460–1478. 10.1161/CIRCRESAHA.118.310082.

19. Griffiths, E.J., and Rutter, G.A. (2009). Mitochondrial calcium as a key regulator of mitochondrial ATP production in mammalian cells. Biochim. Biophys. Acta - Bioenerg. 1787, 1324–1333. 10.1016/j.bbabio.2009.01.019.

20. Hoffman, N.E., Chandramoorthy, H.C., Shanmughapriya, S., Zhang, X.Q., Vallem, S., Doonan, P.J., Malliankaraman, K., Guo, S., Rajan, S., Elrod, J.W., et al. (2014). SLC25A23 augments mitochondrial Ca2+ uptake, interacts with MCU, and induces oxidative stress-mediated cell death. Mol. Biol. Cell 25, 936–947. 10.1091/mbc.E13-08-0502.

21. Guerrero, A., Herranz, N., Sun, B., Wagner, V., Gallage, S., Guiho, R., Wolter, K., Pombo, J., Irvine, E.E., Innes, A.J., et al. (2019). Cardiac glycosides are broad-spectrum senolytics. Nat. Metab. 1, 1074–1088. 10.1038/s42255-019-0122-z.

22. Ahumada-Castro, U., Puebla-Huerta, A., Cuevas-Espinoza, V., Lovy, A., and Cardenas, J.C. (2021). Keeping zombies alive: The ER-mitochondria Ca2+ transfer in cellular senescence. Biochim. Biophys. Acta - Mol. Cell Res. 1868, 119099. 10.1016/j.bbamcr.2021.119099.

23. Martin, N., Zhu, K., Czarnecka-Herok, J., Vernier, M., and Bernard, D. (2023). Regulation and role of calcium in cellular senescence. Cell Calcium 110, 102701. 10.1016/j.ceca.2023.102701.

24. Triana-Martínez, F., Picallos-Rabina, P., Da Silva-Álvarez, S., Pietrocola, F., Llanos, S., Rodilla, V., Soprano, E., Pedrosa, P., Ferreirós, A., Barradas, M., et al. (2019). Identification and characterization of Cardiac Glycosides as senolytic compounds. Nat. Commun. 10. 10.1038/s41467-019-12888-x.

25. Brookes, P.S., Yoon, Y., Robotham, J.L., Anders, M.W., and Sheu, S.-S. (2004). Calcium, ATP, and ROS: a mitochondrial love-hate triangle. Am. J. Physiol. Cell Physiol. 287, C817–33. 10.1152/ajpcell.00139.2004.

26. Krebs, J., Agellon, L.B., and Michalak, M. (2015). Ca(2+) homeostasis and endoplasmic reticulum (ER) stress: An integrated view of calcium signaling. Biochem. Biophys. Res. Commun. 460, 114–121. 10.1016/j.bbrc.2015.02.004.

27. Zhang, Y., Chen, X., Gueydan, C., and Han, J. (2018). Plasma membrane changes during programmed cell deaths. Cell Res. 28, 9–21. 10.1038/cr.2017.133.

28. Bjørkøy, G., Lamark, T., Pankiv, S., Øvervatn, A., Brech, A., and Johansen, T. (2009). Monitoring autophagic degradation of p62/SQSTM1. Methods Enzymol. 452, 181–197. 10.1016/S0076-6879(08)03612-4.

29. Feng, X., Du, W., Ding, M., Zhao, W., Xirefu, X., Ma, M., Zhuang, Y., Fu, X., Shen, J., Zhang, J., et al. (2022). Myosin 1D and the branched actin network control the condensation of p62 bodies. Cell Res. 32, 659–669. 10.1038/s41422-022-00662-6.

30. Hung, W.Y., Chang, J.H., Cheng, Y., Cheng, G.Z., Huang, H.C., Hsiao, M., Chung, C.L., Lee, W.J., and Chien, M.H. (2019). Autophagosome accumulation-mediated ATP energy deprivation induced by penfluridol triggers nonapoptotic cell death of lung cancer via activating unfolded protein response. Cell Death Dis. 10. 10.1038/s41419-019-1785-9.

31. Liu, Y., Hao, Y., Li, Y., Zheng, Y., Dai, J., Zhong, F., Wei, W., and Fang, Z. (2020). Salinomycin induces autophagic cell death in salinomycin-sensitive melanoma cells through inhibition of autophagic flux. Sci. Rep. 10, 1–14. 10.1038/s41598-020-75598-1.

32. Wang, L., Leite de Oliveira, R., Wang, C., Fernandes Neto, J.M., Mainardi, S., Evers, B., Lieftink, C., Morris, B., Jochems, F., Willemsen, L., et al. (2017). High-Throughput Functional Genetic and Compound Screens Identify Targets for Senescence Induction in Cancer. Cell Rep. 21, 773–783. 10.1016/j.celrep.2017.09.085.

33. Takeda, K., Yamaguchi, N., Akiba, H., Kojima, Y., Hayakawa, Y., Tanner, J.E., Sayers, T.J., Seki, N., Okumura, K., Yagita, H., et al. (2004). Induction of tumor-specific T cell immunity by anti-DR5 antibody therapy. J. Exp. Med. 199, 437–448. 10.1084/jem.20031457.

34. Domen, A., Deben, C., De Pauw, I., Hermans, C., Lambrechts, H., Verswyvel, J., Siozopoulou, V., Pauwels, P., Demaria, M., van de Wiel, M., et al. (2022). Prognostic implications of cellular senescence in resected non-small cell lung cancer. Transl. lung cancer Res. 11, 1526–1539. 10.21037/tlcr-22-192.

35. Zhai, J., Han, J., Li, C., Lv, D., Ma, F., and Xu, B. (2023). Tumor senescence leads to poor survival and therapeutic resistance in human breast cancer. Front. Oncol. 13, 1–16. 10.3389/fonc.2023.1097513.

36. Sanjana, N.E., Shalem, O., and Zhang, F. (2014). Improved vectors and genome-wide libraries for CRISPR screening, 10.1038/nmeth.3047 10.1038/nmeth.3047.

37. Doench, J.G., Fusi, N., Sullender, M., Hegde, M., Vaimberg, E.W., Donovan, K.F., Smith, I., Tothova, Z., Wilen, C., Orchard, R., et al. (2016). Optimized sgRNA design to maximize activity and minimize off-target effects of CRISPR-Cas9. Nat. Biotechnol. 34, 184–191. 10.1038/nbt.3437.

38. Evers, B., Jastrzebski, K., Heijmans, J.P.M., Grernrum, W., Beijersbergen, R.L., and Bernards, R. (2016). CRISPR knockout screening outperforms shRNA and CRISPRi in identifying essential genes. Nat. Biotechnol. 34, 631–633. 10.1038/nbt.3536.

39. Sturm, M., Schroeder, C., and Bauer, P. (2016). SeqPurge: Highly-sensitive adapter trimming for paired-end NGS data. BMC Bioinformatics 17, 1–7. 10.1186/s12859-016-1069-7.

40. Kim, D., Paggi, J.M., Park, C., Bennett, C., and Salzberg, S.L. (2019). Graph-based genome alignment and genotyping with HISAT2 and HISAT-genotype. Nat. Biotechnol. 37, 907–915. 10.1038/s41587-019-0201-4.

41. Howe, K.L., Achuthan, P., Allen, J., Allen, J., Alvarez-Jarreta, J., Ridwan Amode, M., Armean, I.M., Azov, A.G., Bennett, R., Bhai, J., et al. (2021). Ensembl 2021. Nucleic Acids Res. 49, D884–D891. 10.1093/nar/gkaa942.

42. He, S., et al. (2024). "Dasabuvir alleviates 5-fluorouracil-induced intestinal injury through anti-senescence and anti-inflammatory." Scientific Reports 14(1).

43. Sun, T., et al. (2022). "Characterization of cellular senescence in doxorubicin-induced aging mice." Experimental Gerontology 163: 111800.

44. Xia, J., et al. (2022). "Metformin ameliorates 5-fluorouracil-induced intestinal injury by inhibiting cellular senescence, inflammation, and oxidative stress." International Immunopharmacology 113: 109342.

45. Gupta, P. B., et al. (2009). "Identification of Selective Inhibitors of Cancer Stem Cells by High-Throughput Screening." Cell 138(4): 645–659.

