## Supplemental figures Casagrande for "An antibiotic that mediates immune destruction of senescent cancer cells"

### AI appendix Fig. S1.

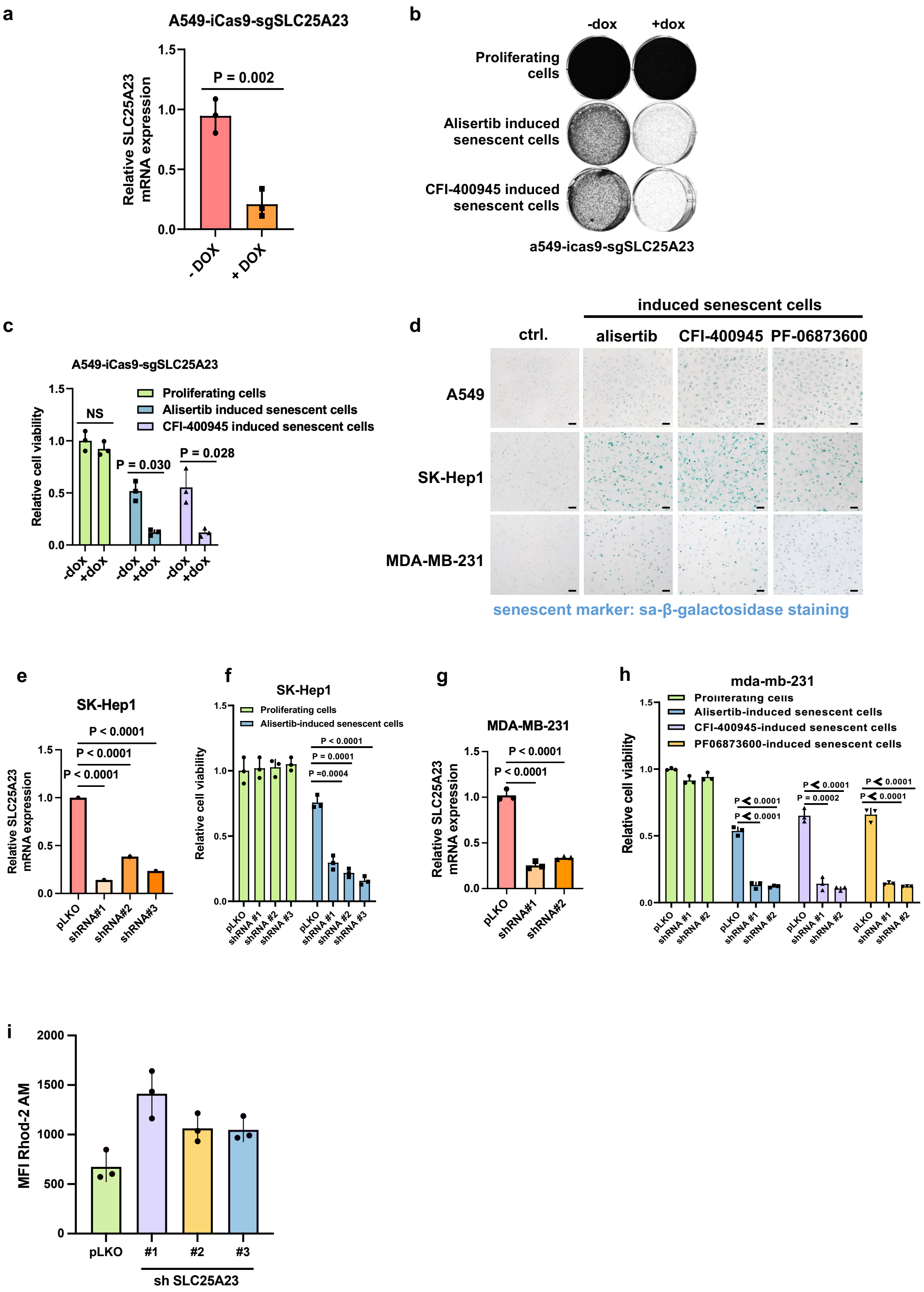

**AI appendix Fig. S1: SLC25A23 is a vulnerability in multiple senescent tumor models**

(a). Real-time PCR analysis of SLC25A23 in A549-iCas9 cells with shRNA against SLC25A23 treated with 1 µg $ml^{-1}$ DOX for 4 days. GAPDH served as a loading control. n = 3 independent experiments. Error bars represent the mean ± standard deviation. Two-Tailed unpaired student's t-test was performed to calculate statistical significance.

(b-c). Cell viability of proliferating and alisertib- and CFI-400945-induced senescent A549-iCas9 with sgRNA against SLC25A23 treated with 1 µg $ml^{-1}$ DOX for 10 days was assessed by Colony Formation Assay. Untreated cells were taken as a control. Cell viability is quantified. n = 3 independent experiments. Error bars represent the mean ± standard deviation. Two-Way ANOVA was performed to calculate statistical significance.

(d). A549, SK-Hep1 and MDA-MB-231 cells were stained with senescence marker SA-β-galactosidase staining to prove senescence induction after 7 days of 0.5 µM alisertib, 100 nM CFI-400945 or 0.5 µM PF-06873600 treatment. Untreated cells were taken as a control.

(e). Real-time PCR analysis of SK-Hep1 cells with independent shRNAs against *SLC25A23* to validate the knockdown efficiency. pLKO empty vector were taken as a control. n = 2 individual replicates. Error bars represent standard deviation. One-Way ANOVA was performed to calculate statistical significance.

(f). Cell viability of proliferating and alisertib-induced senescent SK-Hep1 cells with three independent shRNAs against *SLC25A23* was assessed by Colony Formation Assay. pLKO empty vector was taken as a control.

(g). Real-time PCR analysis of MDA-MB-231 cells with independent shRNAs against *SLC25A23* to validate the knockdown efficiency. pLKO empty vector was taken as a control. n = 3 individual replicates. Error bars represent standard deviation. One-Way ANOVA was performed to calculate statistical significance.

(h). Cell viability of proliferating and alisertib-, CFI-400945-, and PF-06873600-induced senescent MDA-MB-231 cells with two independent shRNAs against *SLC25A23* was assessed by Colony Formation Assay. pLKO empty vector was taken as a control. Cell viability is quantified. n = 3 independent experiments. Error bars represent the mean ± standard deviation. Two-Way ANOVA was performed to calculate statistical significance.

(i). Flow cytometry was used to determine the intensity of $Ca^{2+}$ dye Rhod-2 in A549 cells with three independent shRNAs against SLC25A23. n = 3 individual replicates. Error bars represent the mean ± standard deviation. One-Way ANOVA was performed to calculate statistical significance. Knockdown efficiency was validated with western blot. pLKO was used as a non-targeting shRNA for control. GAPDH served as a loading control.

### AI appendix Fig. S2.

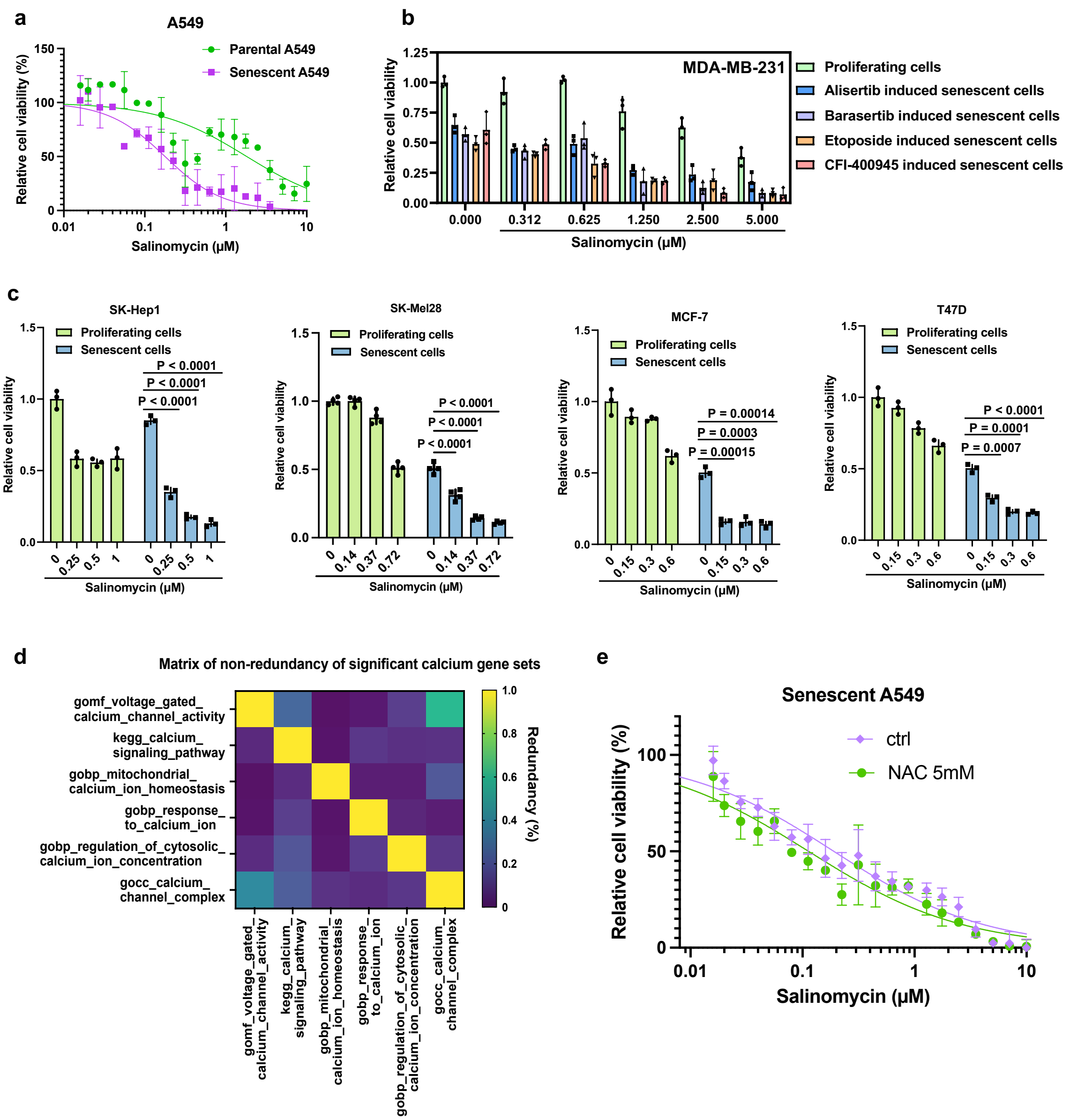

**AI appendix Fig. S2: Salinomycin acts as a senolytic drug in multiple models**

(a). Dose-response curve showing cell viability of alisertib-induced senescent and parental A549 cells after 7 days of salinomycin treatment. Cell viability was assessed by CellTiter-Blue assay. Data was quantified by calculating the IC50 value. The dose-response curves are a significant representation of n = 3 independent experiments. Error bars represent the mean ± standard deviation. Two-tailed unpaired student's t-test was performed to calculate statistical significance.

(b). Cell viability of alisertib-, barasertib-, etoposide- and CFI-400945-induced senescent MDA-MB-231 cells after 5 days of salinomycin treatment was assessed by colony formation assay. Proliferating cells were taken as a control. n = 3 independent experiments. Error bars represent mean + standard deviation. Statistical significance was calculated with Two-Way ANOVA, but is not shown in the graph.

(c). Cell viability of proliferating and alisertib-induced senescent SK-Hep1, SK-Mel28, MCF-7 and T47D cells after 7 days of salinomycin treatment was assessed by CellTiter-Blue assay. Untreated cells were taken as a control. n = 3 independent experiments. Error bars represent mean ± standard deviation. Two-Way ANOVA was performed to calculate statistical significance.

(d). The matrix shows the percentage of overlap of the genes represented in the different selected calcium gene signatures. Values are normalized and plotted between 1 for complete overlap of the signatures and 0 for no overlap.

(e). Cell viability of alisertib-induced senescent A549 cells after 7 days of salinomycin treatment in combination with 5 mM ROS scavenger NAC was assessed by CellTiter-Blue assay to generate the dose-response curve, n = 3 individual replicates. Error bars represent the mean ± standard deviation.

### AI appendix Fig. S3

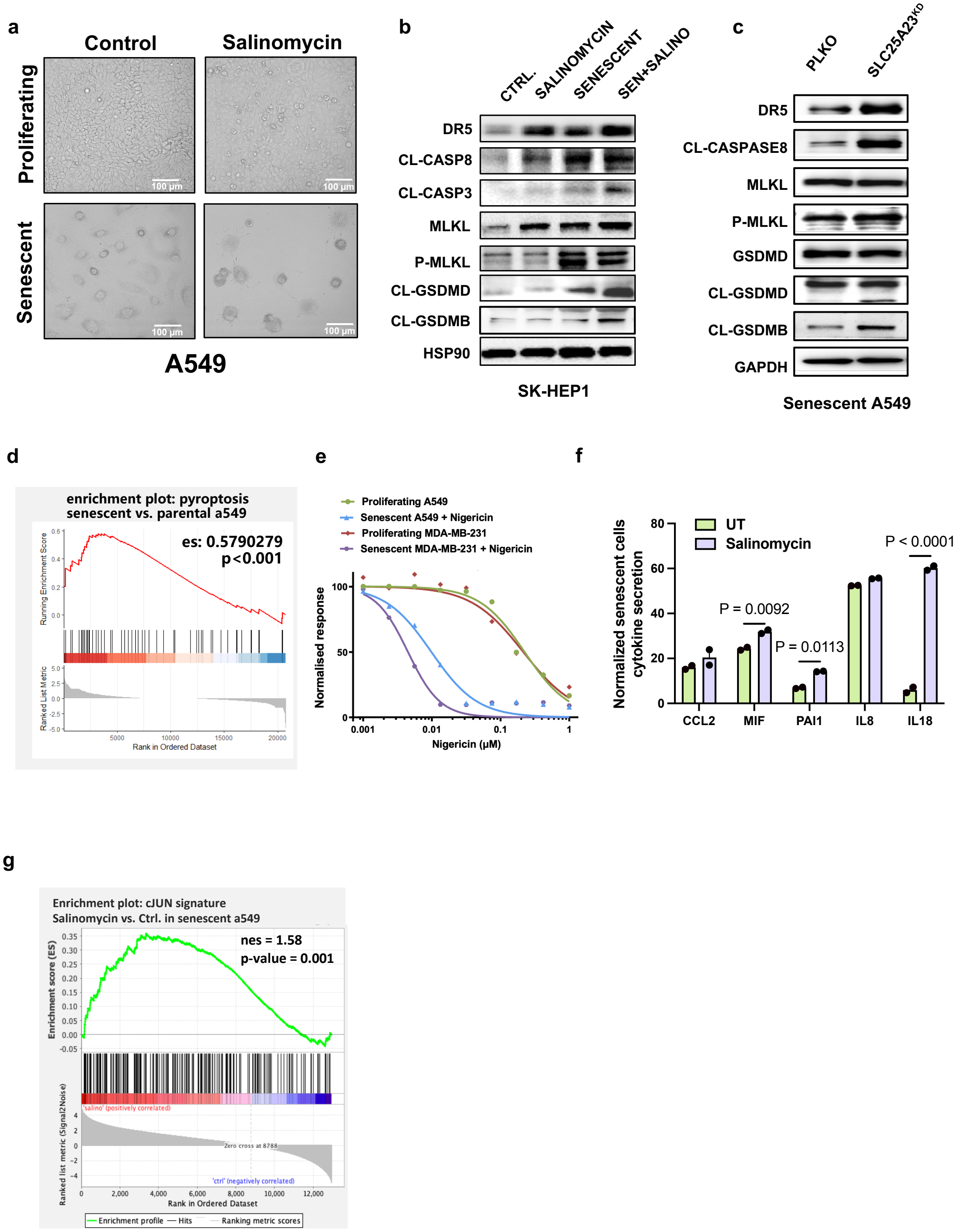

**AI appendix Fig. S3: Salinomycin induces cell death through multiple parallel mechanisms in various models**

(a). Phase contrast images of Salinomycin treated senescent and parental A549. Salinomycin concentration used was 2.5 µM for 72 hours.

(b). Western blot analysis of apoptosis, pyroptosis, necroptosis and autophagy death pathway markers in proliferating and alisertib-induced senescent SK-Hep1 cells after 48 hours of 2.5 µM salinomycin treatment. Untreated proliferating cells were taken as a control. GAPDH served as a loading control.

(c). Western blot analysis of apoptosis, pyroptosis, necroptosis and autophagy death pathway markers in alisertib-induced senescent A549 cells with shRNA against SLC25A23. GAPDH served as a loading control.

(d). GSEA signature for pyroptosis of senescent cells vs. proliferating A549.

(e). Dose-response curve showing cell viability of proliferating and alisertib-induced senescent A549 and MDA-MB-231 cells after 7 days of nigericin treatment. Cell viability was assessed by CellTiter-Blue assay.

(e). Quantification of differentially expressed cytokines between salinomycin treated and ctrl senescent A549 cells. A cytokine array was performed to measure the secretion of alisertib-induced senescent A549 cells after 72 hours of 2.5 µM salinomycin treatment. Untreated senescent cells were taken as a control.

(f). CREP1-cJun gene signature enrichment analysis (GSEA) on transcriptomic data in alisertib-induced senescent A549 cells compared to proliferating cells.

### AI appendix Fig. S4

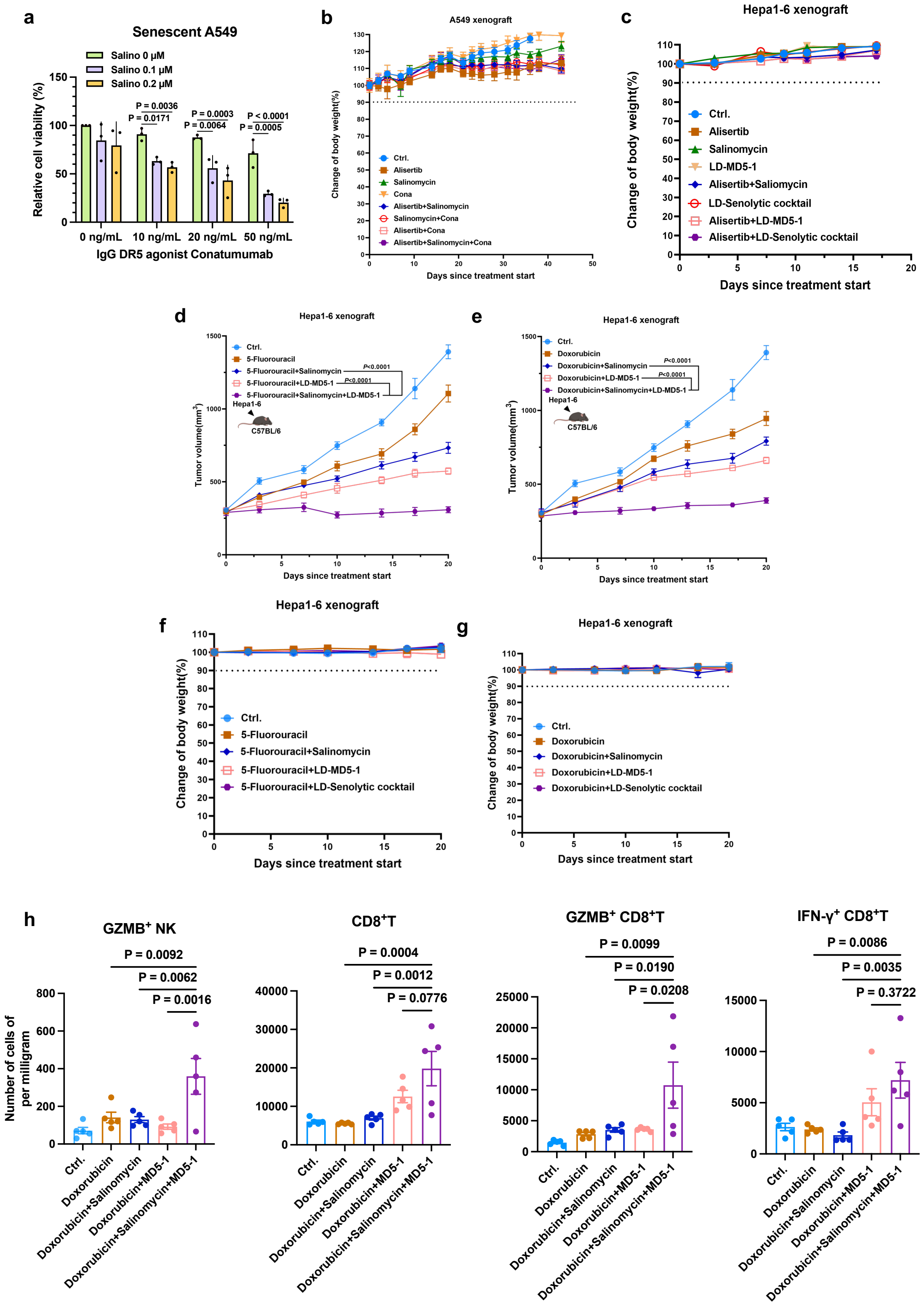

**AI appendix Fig. S4: cJUN is activated in salinomycin treated senescent cells, and induces the DR5 upregulation**

(a). Cell viability of alisertib-induced senescent A549 treated with 0, 10, 20 or 50 ng $ml^{-1}$ DR5 agonist conatumumab in combination with 0, 0.1 or 0.2 μM salinomycin was assessed by CellTitel-Blue Assay. n = 3 independent experiments. Error bars represent mean + standard deviation. Two-Way ANOVA was performed to calculate statistical significance.

(b). Body weight measurement of mice tracked throughout the duration of the experiment. When tumors reached approximately 100 $mm^3$, animals were assigned to either vehicle control, 25 mg kg-1 alisertib, 1.5 mg kg-1 salinomycin, 5 μg per dose conatumumab (Cona), the two-way combinations of 1.5 mg kg-1 salinomycin plus 5 μg per dose conatumumab and 25 mg kg-1 alisertib plus 5 μg per dose conatumumab, or the three-way combination of 25 mg kg-1 alisertib plus 1.5 mg kg-1 salinomycin plus 2.5 μg per dose conatumumab.

(c, f-g). Body weight of mice experiment with Hepa1-6 murine hepatoma cells into immunocompetent C57BL/6 mice for (c), n = 6 mice per group. For (f-g), n = 5 mice per group.

(d-e). Tumor growth of Hepa1-6 murine hepatoma cells in syngeneic immunocompetent C57BL/6 mice. When tumors reached approximately 250 $mm^3$, animals were assigned to either vehicle control or 40 mg kg-1 5-fluorouracil (d) and 3 mg kg−1 doxorubicin (e) as senescent inducers. The senolytic cocktail dosage was constant for the three experiments, equal to 1.5 mg kg-1 salinomycin, 0.3 μg per dose MD5-1, used as single or double agents and in a three-way combination with the different senescence inducers. For (d-e), n = 5 mice per group

(h). Quantitative estimate of various immune effector cells per milligram of tumor tissue in the tumors with different treatment groups, as determined by flow cytometry. n = 5 tumors per group.

### AI appendix Fig. S5

a

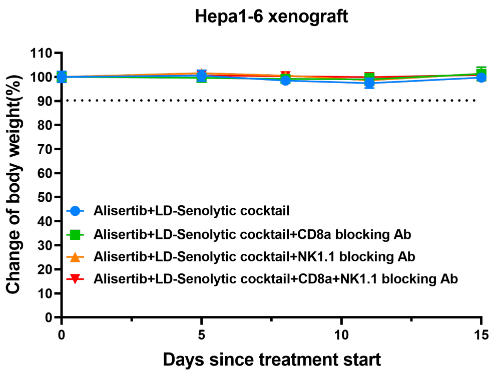


b

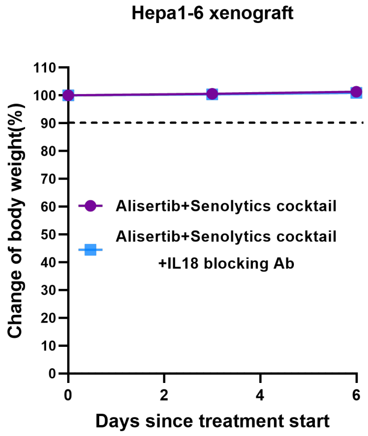

**AI appendix Fig. S5: Body weight of mice treated with the senolytic cocktail indicates no major toxicity**

(a-b). Body weight of mice experiment with Hepa1-6 murine hepatoma cells into immunocompetent C57BL/6 mice for (a), n = 5 mice per group. For (b), n = 5 mice per group.

### Supplementary Table 1

| Drug Name | Function | Category |
|---|---|---|
| Fccp | OXPHOS uncoupler | OXPHOS |
| Pyrvinum pamoate | OXPHOS inhibitor | OXPHOS |
| antimycin A | Complex III inhibitor | Mitochondrial complex |
| atovaquone | Complex III inhibitor | Mitochondrial complex |
| fenofibrate | Complex I inhibitor | Mitochondrial complex |
| devimistat | Complex I inhibitor | Mitochondrial complex |
| mitoxantrone | Complex I inhibitor | Mitochondrial complex |
| celastrol | Complex I inhibitor | Mitochondrial complex |
| elesclomol | Complex I inhibitor | Mitochondrial complex |
| vlx600 | Complex I inhibitor | Mitochondrial complex |
| bay-872243 | Complex I inhibitor | Mitochondrial complex |
| metformin | Complex I inhibitor | Mitochondrial complex |
| canagliflozin | Complex I inhibitor | Mitochondrial complex |
| rotenone | Complex I inhibitor | Mitochondrial complex |
| Tigecyclin | mitoribosome disruptor | Mitochondrial funcion |
| Doxycycline | mitoribosome disruptor | Mitochondrial funcion |
| mupirocin | mitochondrial isoleucyl tRNA synthetase inhibitor | Mitochondrial funcion |
| docetaxel | mitochondrial disruptor | Mitochondrial funcion |
| cisplatin | mitochondrial disruptor | Mitochondrial funcion |
| Actinonin | mitochondrial deformylase inhibitor | Mitochondrial funcion |
| thapsigargin | SERCA inhibitor | ER stress modulatory |
| 2abp | IP3R1 antagonist | ER stress modulatory |
| carbachiol | IP3R1 agonist | ER stress modulatory |
| Chloroquine | Autophagy inhibitor | ER stress modulatory |
| ru360 | MCU inhibitor | Calcium regulatory |
| benzethonium | MCU antagonist | Calcium regulatory |
| amorolfine | MCU agonist | Calcium regulatory |
| Ionomycin | Calcium ionophore | Calcium regulatory |
| Salinomycin | Calcium ionophore | Calcium regulatory |
| BAPTA-AM | Calcium chelator | Calcium regulatory |
| amlodipine | calcium channel blocker | Calcium regulatory |
| cinacalcet | calcium channel blocker | Calcium regulatory |
| furosemide | calcium channel blocker | Calcium regulatory |
| bay K 8644 | calcium channel blocker | Calcium regulatory |
| Diltiazem | calcium channel blocker | Calcium regulatory |
| Kn90 | calcium channel blocker | Calcium regulatory |
| Nifedipine | calcium antagonist | Calcium regulatory |
| Berbamine | calcium antagonist | Calcium regulatory |
| Verapamil | calcium antagonist | Calcium regulatory |
| abt 263 | BCL2 family inhibitor | Positive Control |
